# Structure and functional analysis of the *Legionella* chitinase ChiA reveals a novel mechanism of metal-dependent mucin degradation

**DOI:** 10.1101/687871

**Authors:** Katherine H. Richardson, Lubov S. Grigoryeva, Paula Corsini, Richard C. White, Rosie Shaw, Theo J. Portlock, Benjamin Dorgan, Arianna Fornili, Nicholas P. Cianciotto, James A. Garnett

**Author notes:** Correspondence to: James Garnett, Centre for Host Microbiome Interactions, Dental institute, King’s College London, London, UK., Nicholas Cianciotto, Department of Microbiology and Immunology, Northwestern University Feinberg School of Medicine, Chicago, Illinois, USA.

## Abstract

Chitinases are important enzymes that contribute to the generation of carbon and nitrogen from chitin, a long chain polymer of N-acetylglucosamine that is abundant in insects, fungi, invertebrates and fish. Although mammals do not produce chitin, chitinases have been identified in bacteria that are key virulence factors in severe respiratory, gastrointestinal and urinary diseases. However, it is unclear how these enzymes are able to carry out this dual function. *Legionella pneumophila* is the causative agent of Legionnaires’ disease, an often-fatal pneumonia and its chitinase ChiA is essential for the survival of *L. pneumophila* in the lung. Here we report the first atomic resolution insight into the pathogenic mechanism of a bacterial chitinase. We derive an experimental model of intact ChiA and show how its N-terminal region targets ChiA to the bacterial surface after its secretion. We provide the first evidence that *L. pneumophila* can bind mucins on its surface but this is not dependent on *chiA*. This demonstrates that additional peripheral mucin binding proteins are also expressed in *L. pneumophila*. Finally, we show that the ChiA C-terminal chitinase domain has novel metal-dependent peptidase activity against mammalian mucins. These findings suggest that ChiA facilitates bacterial penetration of the alveolar mucosa and ChiA may be a promising target for vaccine development.

## Introduction

*Legionella pneumophila* is a Gram-negative bacterium that can withstand large variation in pH and temperature. When humans are exposed to *L. pneumophila*, it can infect macrophages and epithelia in the lungs and trigger chronic inflammation and tissue damage (1). *L. pneumophila* is the causative agent of Legionnaires’ disease, an often-fatal pneumonia, and Pontiac fever, a milder flu-like disease, and rates of infection are increasing each year (1–4). Although infection is primarily via inhalation of contaminated water droplets from aerosolizing devices (5), there is also now evidence for person-to-person transmission (6, 7).

Upon invasion of eukaryotic hosts, *L. pneumophila* avoids fusion with canonical endosomal/lysosomal pathways by forming a membrane bound compartment, the *Legionella* containing vacuole (LCV) (1). *L. pneumophila* encodes over 300 proteins that are exported from this modified phagosome into the host cytoplasm by the Icm/Dot type IVb secretion system (1). These effectors manipulate host signalling pathways and mediate evasion of the host’s degradative lysosomal pathway, enabling *L. pneumophila* to replicate to large numbers (8). *L. pneumophila* also expresses a type II secretion system (T2SS), which secretes at least 25 proteins, including almost 20 enzymes and substrates that contain a high proportion of unique amino acid sequence with no homology outside of the *Legionella* genus (9). The T2SS is important for both intracellular and extracellular lifestyles (9). These processes include extracellular growth at low temperatures, biofilm formation, intracellular replication in amoebae and macrophages, dampening of cytokine output from infected cells and persistence in lungs (9–15).

Among the type II substrates, ChiA is an 81 kDa endochitinase with a novel amino acid sequence at its N-terminus and a putative glycosyl hydrolase 18 (GH18) domain at its C-terminus (15). Chitin is an insoluble carbohydrate composed of linear β-1,4-linked N-acetylglucosamine (GlcNAc) residues and its degradation by chitinase enzymes serves as an important source of nutrients for many bacteria (16). Chitin is not synthesized by mammals and *L. pneumophila chiA* mutants are not impaired for growth in *Vermamoeba vermiformis* amoebae and macrophage-like U937 cells (15). However, they are less able to survive in the lungs of A/J mice, suggesting that ChiA is required for optimal survival of *L. pneumophila* in the lungs (15), although how it is able to promote infection is unknown.

In this study, we report a structural model for full-length ChiA based on X-ray crystallographic, template based modelling and small angle X-ray scattering (SAXS) data. ChiA is composed of four domains (N1, N2, N3 and CTD, from N-to C-terminus) which have structural homology to those associated with other chitinase enzymes. Using chitin binding and chitinase activity assays, we show that ChiA-N1 is a chitin binding module and we confirm that ChiA-CTD is a glycosyl hydrolase domain. Using binding assays, we show that both the ChiA-N3 domain and eukaryotic mucins can associate with the *L. pneumophila* surface. Lastly, our structural and biochemical studies demonstrate that ChiA-CTD has novel mucinase activity, independent of its chitinase active site. Our work provides novel molecular insight into the virulence mechanism of a bacterial chitinase and suggests that ChiA has a role in liberating nutrients and facilitating *L. pneumophila* penetration of the alveolar mucosa.

## Results

### ChiA is a multi-domain protein

Full-length ChiA from *L. pneumophila* 130b (ChiA-FL; numbered 1-762 for the mature protein; NCBI accession WP_072401826.1) with an N-terminal His_6_-tag was produced in *Escherichia coli* K12 and purified by nickel-affinity and size exclusion chromatography. Despite extensive screening ChiA-FL resisted crystallization and we therefore used bioinformatics analysis to produce a series of subdomain constructs for further characterization (Fig. 1A). While previous examination of the C-terminal domain of ChiA (ChiA-CTD: residues 419-762) had revealed high primary sequence homology to other GH18 chitinase domains (15), the N-terminal region (ChiA-NT; residues 1-417) contains unique primary sequence with no significant homology to any other known protein. Nonetheless, through secondary structure prediction (17) and template based modelling using the Phyre2 (18) and Robetta (19) servers, we identified three putative N-terminal subdomains based on predicted structural similarity with carbohydrate-binding modules (CBMs; ChiA-N1: residues 1-140), fibronectin type-III-like domains (Fn3; ChiA-N2: residues 138-299) and a chitinase A N-terminal domain (ChiN; ChiA-N3: residues 285-417) (Fig. 1A, Table S1).

**Figure 1:**
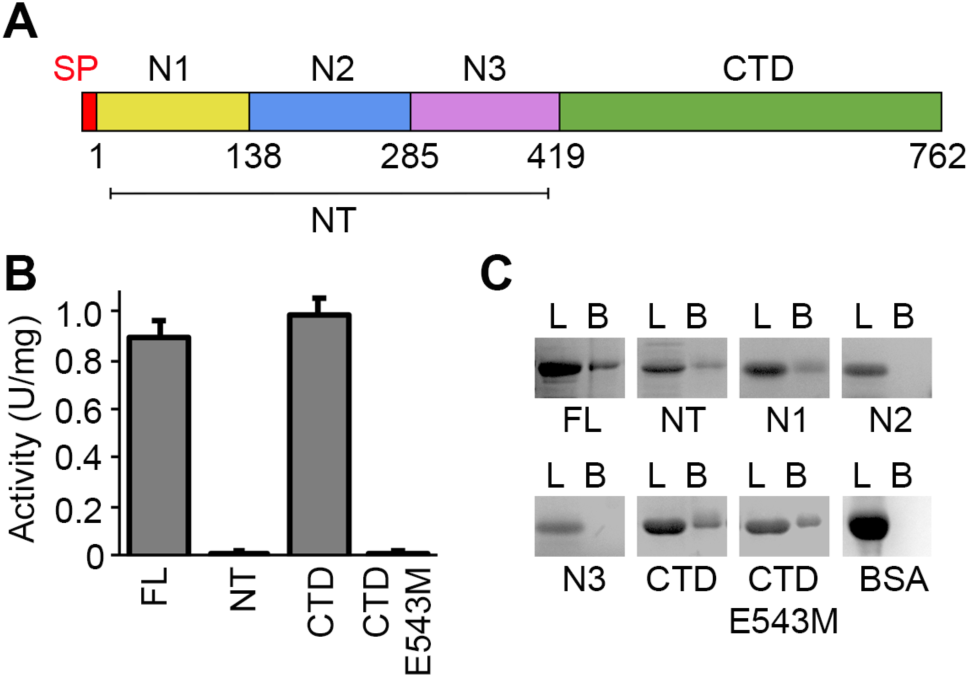
Chitin binding and endochitinase functions of ChiA. (A) Schematic representation of ChiA with domain boundaries annotated. (B) ChiA-FL, subdomains (NT, N1, N2, N3, CTD) and ChiA-CTD^E543M^ were assayed for chitinase activity against *p*-NP-[GlcNAc]_3_. Data represent the mean and standard deviation for triplicate experiments. (C) Chitin pull-down experiment to assess direct interactions between immobilized chitin and ChiA. ChiA-FL and subdomains (NT, N1, N2, N3, CTD, CTD^E543M^) were incubated with chitin beads and analysed by SDS-PAGE. BSA was used as a control. L: loaded sample; B: eluted beads. Data is representative of three independent repeat experiments.

To examine the function of the ChiA N-terminal domains we began by examining their endochitinase activity (15). Each construct was expressed with an N-terminal His_6_-tag in *E. coli* K12 and purified by nickel-affinity and size exclusion chromatography. All reagents were well folded as determined by 1D ^1^H nuclear magnetic resonance (NMR) spectroscopy (Fig. S1). As expected ChiA-FL and ChiA-CTD were both active against *p*-nitrophenyl β-D-*N,N′,N″* triacetylchitotriose (*p*NP-[GlcNAc]_3_) but no activity was detected for ChiA-NT or an E543M ChiA-CTD active site mutant (ChiA-CTD^E543M^) (Fig. 1B, S2). We then assayed binding of ChiA sub-domains to immobilized chitin and observed that in addition to ChiA-CTD, the N1-domain also recognizes chitin polymers, which supports its role as a carbohydrate binding module (Fig. 1C, S3).

**Figure 2:**
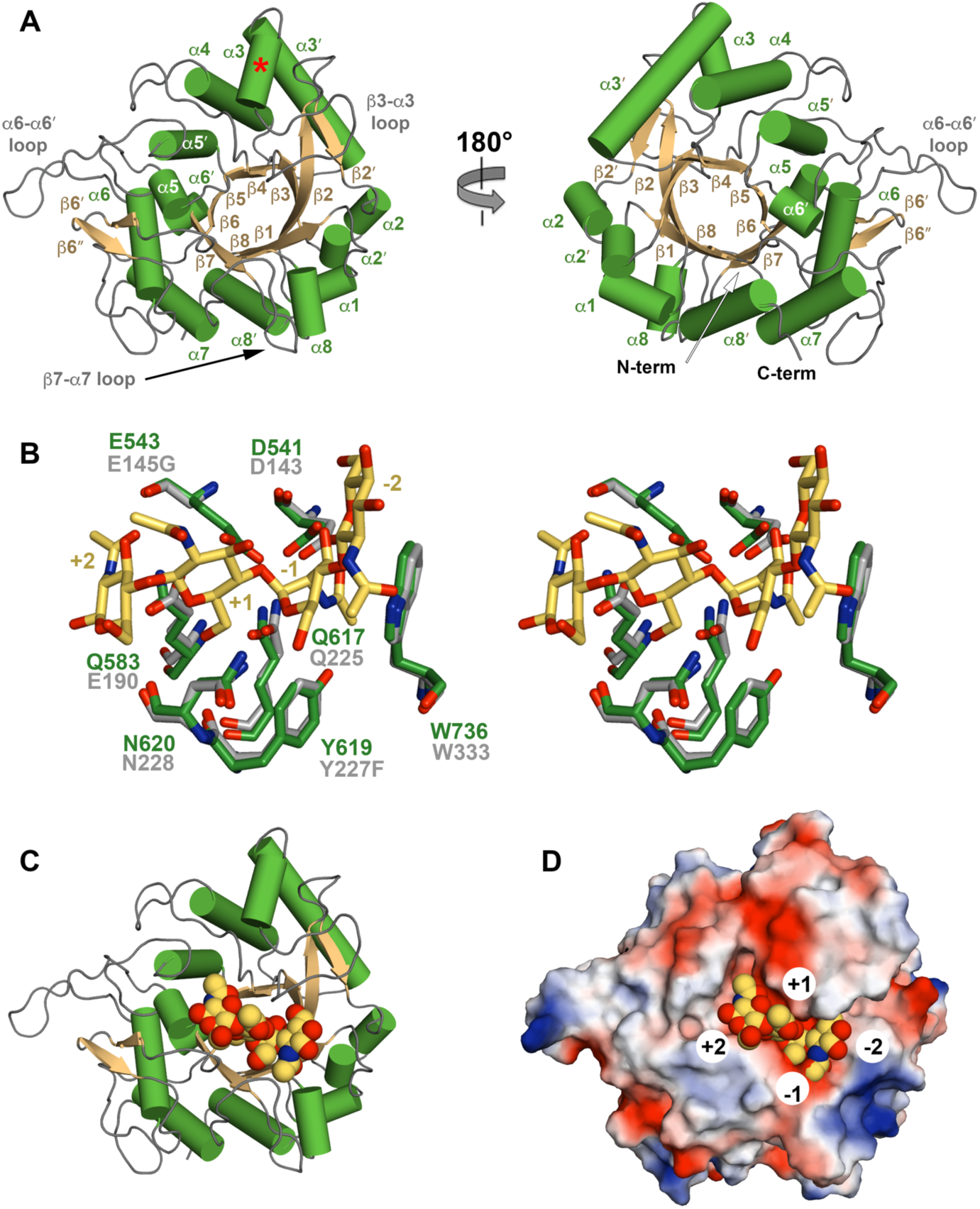
Crystal structure of ChiA-CTD. (A) Cartoon representation of ChiA-CTD with secondary structure and extended loops annotated. Additional ChiA-CTD *α*3-helix is highlighted with a red asterisk. (B) Stick representation of the ChiA-CTD active site (green) superimposed with ChiNCTU2 E145G/Y227F mutant (grey) in complex with chitotetrose (yellow) (PDB ID code 3n18) (21). Mutated residues in ChiNCTU2 are indicated and carbohydrate positions relative to the hydrolysed glycosidic bond are numbered. (C) Model of ChiA-CTD shown as cartoon and (D) electrostatic surface potential, bound to chitotetrose drawn as spheres.

**Figure 3:**
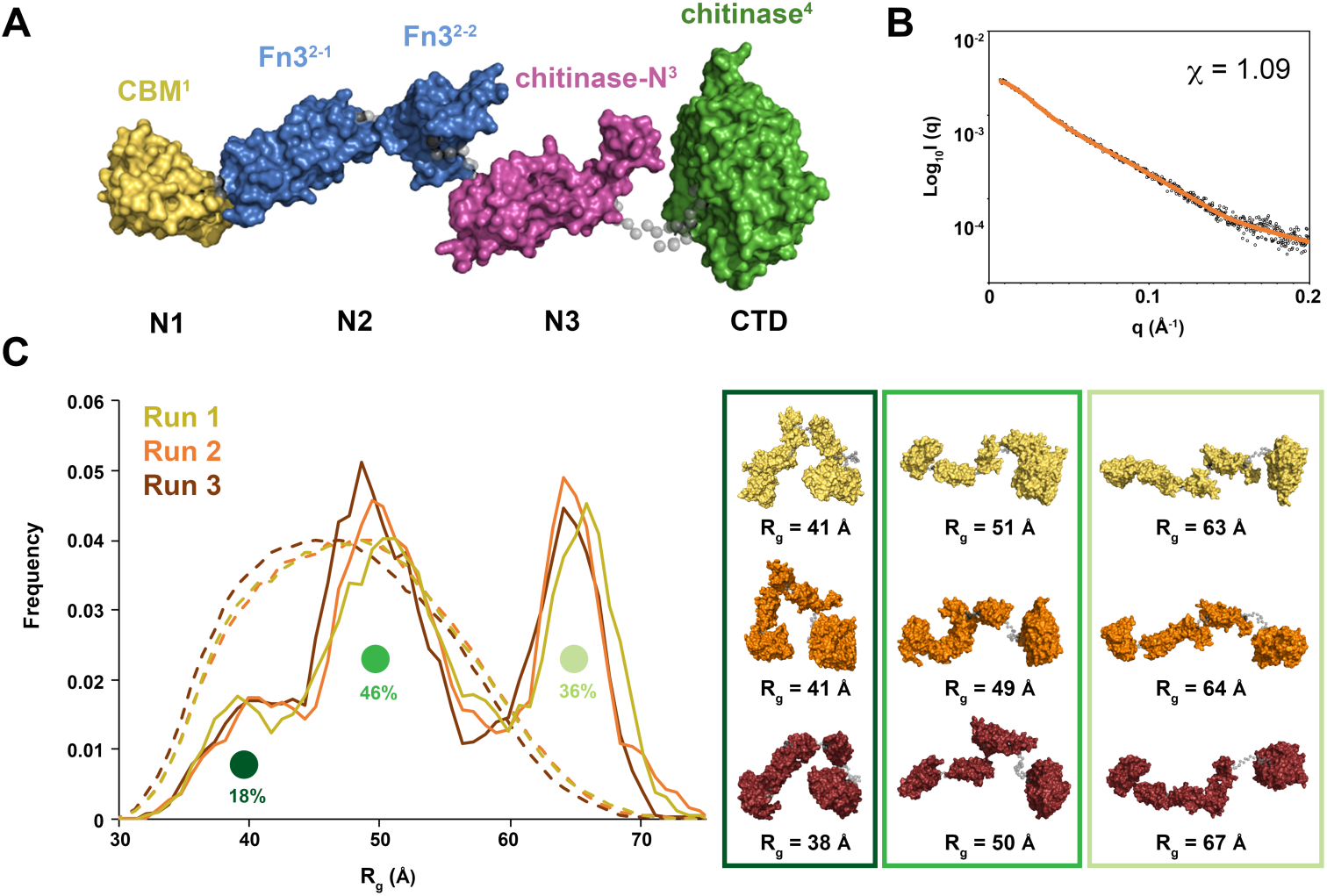
Model of ChiA-FL in solution. (A) Initial model of ChiA generated by EOM 2.0 (24). Linkers are shown as grey spheres. (B) EOM fit (orange line) to the ChiA SAXS data (black open circles) with *χ*^2^ of 1.09. (C) Three independent ensemble optimization method runs (yellow, orange and burgundy) yielded similar distributions of three populations. Sample ChiA models corresponding to the centre of each population for all three runs are shown.

### Atomic structure of ChiA-CTD

We next initiated crystallographic studies of the ChiA subdomains. We readily obtained crystals for ChiA-CTD and its structure was determined using iodide single isomorphous replacement with anomalous scattering (I-SIRAS) phasing. Electron-density maps were refined to 1.7 Å (Table S2) and the final model contains two identical chains, with all molecules built except for the N-terminal His_6_-tags and adjacent ChiA-CTD residues Val419 to Gly424. Each chain forms an anticipated GH18 *α*/*β*-fold and is composed of 11 *β*-strands and 13 *α*-helices (Fig. 2A). High concentrations of 2-methyl-2,4-pentanediol (MPD) were used as a precipitant during crystallization and we observed four MPD molecules in the final model; one bound to the catalytic Asp541 and Glu543 residues (MPD-1) and one within a hydrophobic pocket formed by the *α*5 helix and *α*5-*α*5*′/β*5-*β*6*′* loops (MPD-2) (Fig. S4).

The overall structure of ChiA-CTD is highly similar to other GH18 chitinase domains, and the Dali server (20) identified *Bacillus cereus* ChiNCTU2 enzyme inactive E145G/Y227F mutant in complex with chitotetrose (Protein Data Bank (PDB) ID code 3n18) (21); *Bacillus anthracis* Chi36 (PDB ID code 5kz6); *Chromobacterium violaceum* ChiA (PDB ID code 4tx8); and *Streptomyces coelicolor* ChiA (PDB ID code 3ebv) as having the highest homologies (Z score: 36.3, 35.9, 34.4, 34.1 respectively; rmsd: 2.2 Å, 2.3 Å, 2.2 Å, 1.8 Å, respectively). The chitinase active sites of ChiNCTU2 and ChiA-CTD have high primary sequence identity and tertiary structure homology (Fig. 2B, S5) and modelling of chitotetrose binding indicates that chitin lines a negatively charged valley on the surface of ChiA-CTD. Chitotetrose overlays with MPD-1 in the active site (Fig. 2C, D, S4) and the MPD-2 site is positioned adjacent to the reducing end of the modelled chitotetrose. However, *L. pneumophila* ChiA-CTD also possesses unique features that are not observed in homologous structures. These include an additional *α*-helix (*α*3), an extended *β*3-*α*3 loop, an extended *α*6-*α*6*′* loop and an extended *β*7-*α*7 loop (Fig. 2A, S5, S6).

**Figure 4:**
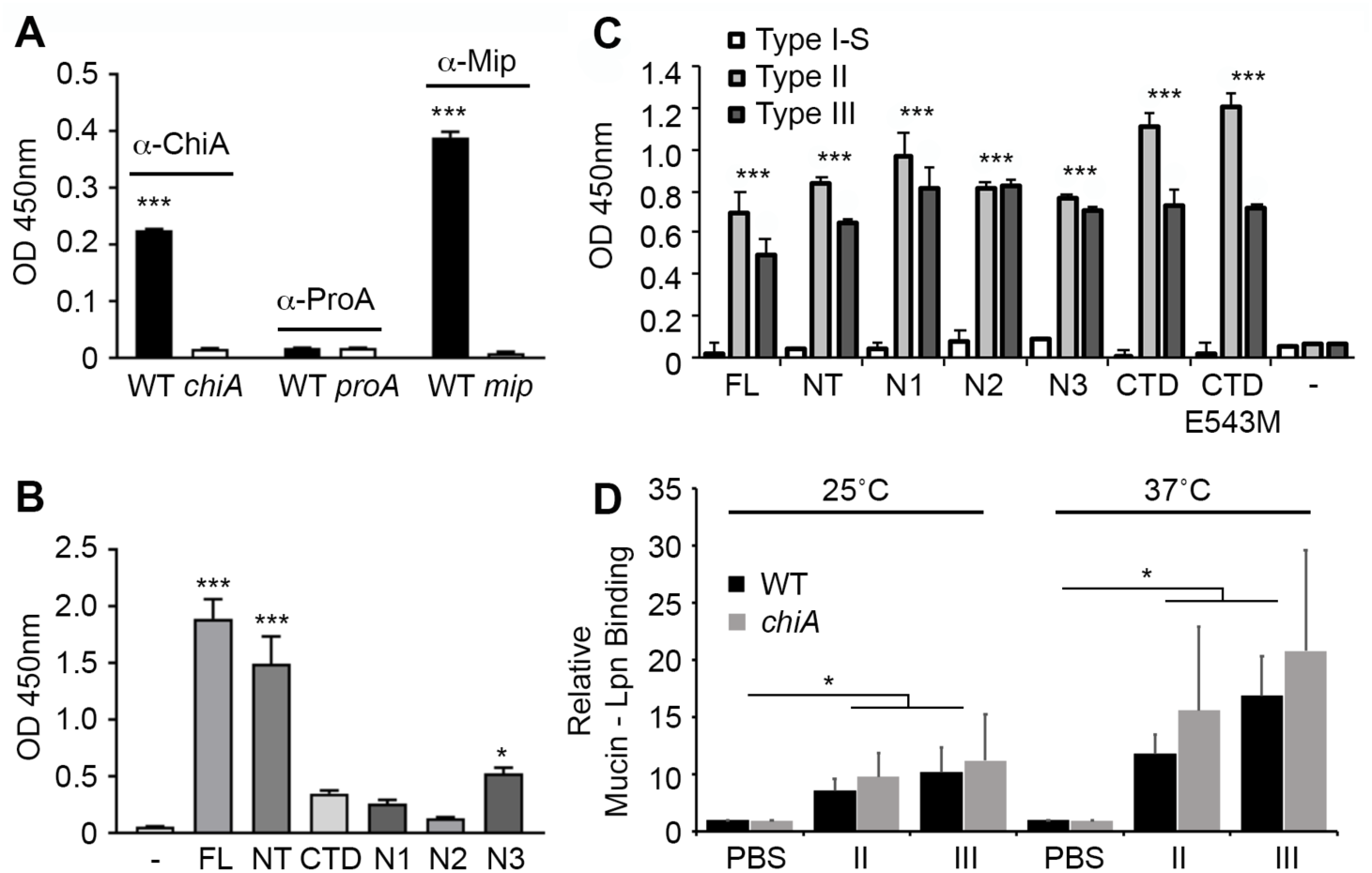
*L. pneumophila* surface association of ChiA and mucin binding. (A) Whole cell ELISA of *L. pneumophila* wild-type 130b (WT), *chiA* mutant NU318 (*chiA*), *proA* mutant AA200 (*proA*) and *mip* mutant NU203 (*mip*) detected with antiserum specific for either ChiA, ProA, or Mip. (B) Whole cell ELISA of *chiA* mutant incubated with either recombinant ChiA-FL or subdomains (NT, N1, N2, N3, CTD) and detected with antiserum specific for either ChiA, ProA, or Mip. PBS buffer alone was used as a control (-). Multiple comparisons against the control were made using a one-way ANOVA, *Holm*-*Šídák* multiple comparisons test; * *P* < 0.05, *** *P* < 0.001. (C) ELISA analysis of binding between immobilised type I-S, II or III porcine stomach mucins and His-tagged ChiA-FL and subdomains (NT, N1, N2, N3, CTD, CTD^E543M^) detected with anti-His-tag antibody. BSA-coated wells were used as controls. *** *P* < 0.001; verses control empty well by two-tailed Student’s test. (D) Mucin binding to GFP-expressing *L. pneumophila* WT or *chiA* mutant strains were incubated at 25 °C or 37°C with PBS, type II or III porcine stomach mucins followed by Texas Red-tagged wheat germ agglutinin (WGA). Mucin binding to bacteria was quantified by flow cytometry. * *P* < 0.05; verses PBS control by two-tailed Student’s test. All data represent the mean and standard deviation for triplicate experiments.

**Figure 5:**
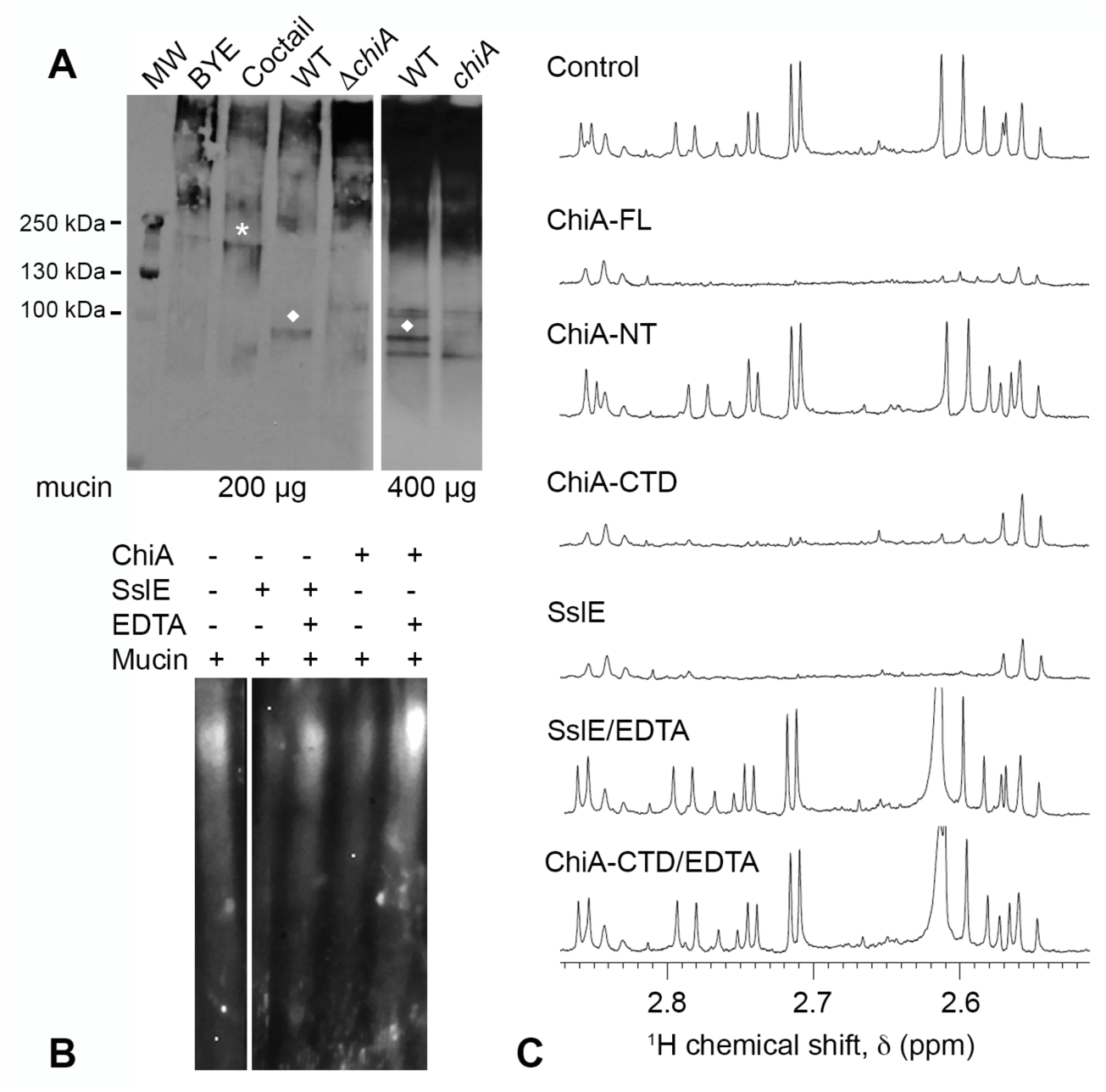
Mucinase activity of ChiA. (A) Secreted mucinase activity of *L. pneumophila* wild-type and *chiA* mutant strains. Immunoblot of type II porcine stomach mucins (200 µg or 400 µg) incubated with either BYE medium alone (BYE), a cocktail of known mucinase enzymes added to BYE medium (cocktail), or supernatants from BYE cultures of wild-type 130b (WT) or *chiA* mutant NU318 (*chiA)*. Asterisk highlights a lower-MW mucin species generated by the cocktail, while diamonds denote an ∼100 kDa mucin species generated by the WT but not mutant supernatants. The data presented are representative of three independent experiments. (B) Immunoblot of type II porcine stomach mucin extract incubated with either ChiA-FL or SslE +/- EDTA or buffer alone. (C) 1D ^1^H NMR spectra between ∼2.5 to ∼2.9 ppm is shown for type II mucin extract incubated with ChiA-FL, ChiA-NT, ChiA-CTD and SslE +/- EDTA or buffer alone.

**Figure 6:**
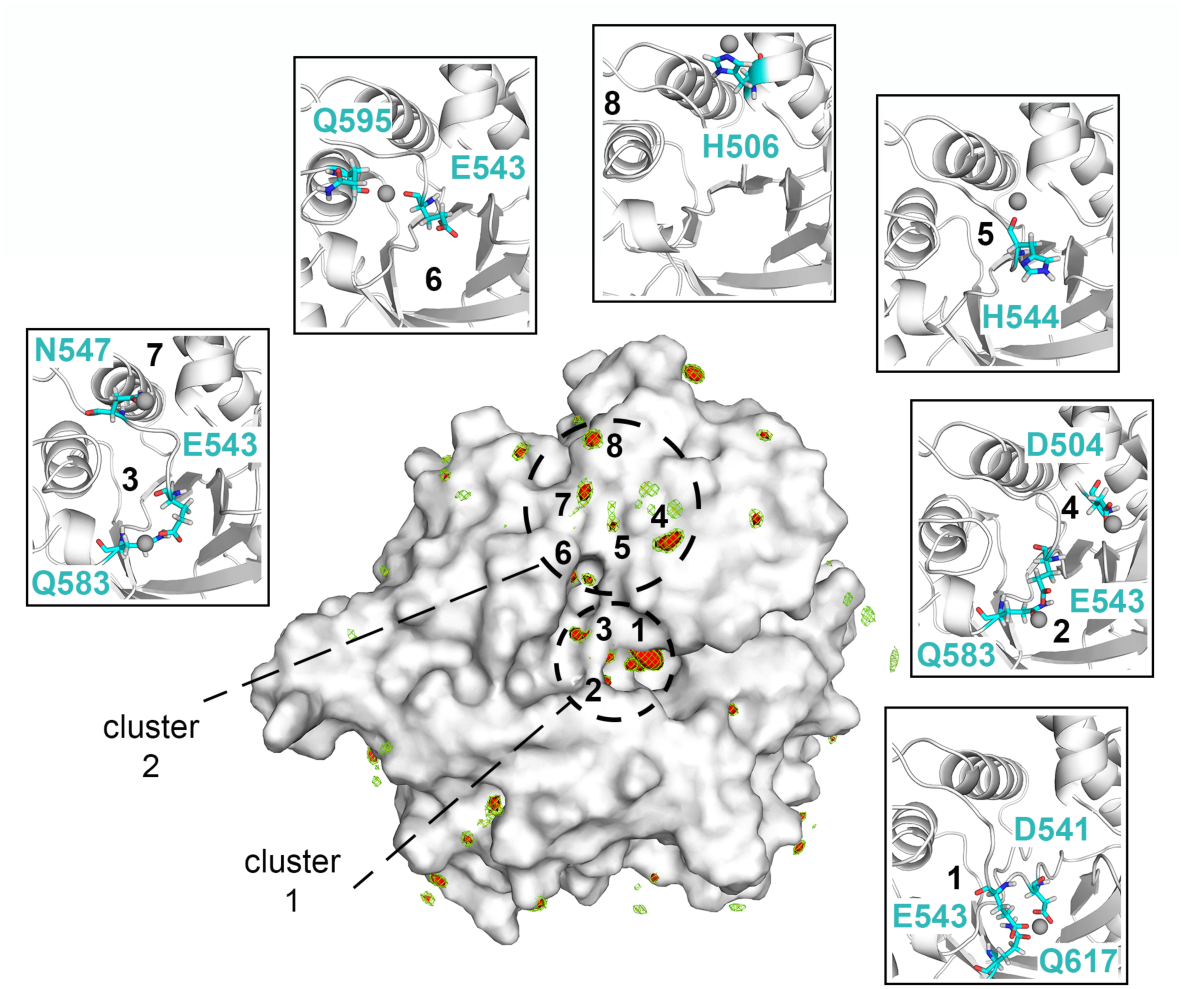
ChiA-CTD Zn^2+^ binding sites. Surface representation of ChiA-CTD showing the spatial distribution of Zn^2+^ ions during MD simulations. The sdf is represented with isosurfaces connecting points with sdf = 20 (green mesh), 25 (yellow mesh) and 30 (red surface) x average sdf. Zn^2+^ high-density sites (red surface) around the chitinase and peptidase active sites are numbered 1 to 8. Blow out boxes show representative structures from the MD simulations to illustrate Zn^2+^ binding in the eight regions, with Zn^2+^ ions shown as spheres, their coordinating residues as sticks and ChiA-CTD as cartoon.

### ChiA is an elongated and dynamic structure in solution

We used small angle X-ray scattering (SAXS) to model the global structure of full-length ChiA in solution. Four different concentrations at 4, 2, 1, and 0.5 mg/ml were measured but signs of aggregation were apparent at concentrations above 1 mg/ml (data not shown). To achieve the highest signal/noise ratio, all further analysis was carried out with the data from the 1 mg/ml sample (Fig. S7, Table S3). Guinier analysis suggested a radius of gyration (*R*_g_), the root mean square distance to the particles centre of mass, of 5.43 nm and analysis of the distance distribution function (P(r)) suggested a maximum particle dimension (*D*_max_) of 17.77 nm and *R*_g_ of 5.45 nm (Fig. S7). Using BSA as a standard, we calculated a particle molecular mass of 89.2 kDa, which is within the method error range for a monomeric 82.6 kDa ChiA.

Kratky, Kratky-Debye and Porod-Debye plot analyses of the SAXS data indicated that ChiA is a highly dynamic particle in solution (Fig. S7) (22). This is likely due to flexibility within the ChiA inter-domain linkers and we therefore used the ensemble optimization method (EOM) to determine molecular model ensembles of ChiA that best fit the SAXS data (23). As we were not able to obtain crystals for ChiA N-domains, an initial model of ChiA was created using a Phyre2 derived N1-domain (residues 22-147), a Robetta derived N2-domain (encompassing two further subdomains: residues 152-245 and 248-305), a Phyre2 derived ChiA N3-domain (residues 315-414) and the crystal structure of ChiA-CTD (residues 439-777), separated by flexible linkers (Fig. 3A; Table S1) (18, 19, 23). Ensemble optimization analysis of the scattering data again supported ChiA being highly flexible in solution (R_flex_ 91.4) (24), with conformation ensembles clustered within three populations (Fig. 3B, C; Table S4). The majority of the simulated conformations exhibited partially extended or fully extended structures at *R_g_* 44-56 Å or *R_g_* 60-71 Å, respectively, whereas minor conformations of closed structures were also populated at *R_g_* 33-43 Å. Inter-domain flexibility is a key feature of processive enzymes (25) and these data suggest that processivity is also important for ChiA function.

### ChiA is targeted to the *L. pneumophila* surface

Although we have repeatedly detected secreted ChiA in bacterial culture supernatants (15, 26), it has been well documented in other bacteria that some type II substrates associate with the bacterial surface upon their secretion (27, 28). To determine whether ChiA is also targeted to the bacterial surface, *L. pneumophila* 130b was examined by whole-cell ELISA using anti-ChiA, anti-Mip and anti-ProA antibodies (Fig. 4A). Both ChiA and Mip, a known surface exposed protein (29), were positive by ELISA, whilst the metalloprotease ProA, another T2SS substrate (9, 30), was negative. To establish which region of ChiA is responsible for this localization, *L. pneumophila* NU318 (*chiA*) mutant was incubated with recombinant ChiA fragments and the whole-cell ELISA was repeated using ChiA antisera (Fig. 4B). ChiA-NT and the N3-domain exhibited significant binding to the bacterial surface while no binding was observed for ChiA-CTD, ChiA-N1 or ChiA-N2.

### ChiA is a mucin binding protein

Some bacterial chitinases and chitin binding proteins are able to promote infection through adhesion to and/or degradation of host glycoconjugates (31) and we hypothesized that ChiA may interact with exogenous mucins in the lungs and elsewhere. We therefore examined the binding capacity of recombinant ChiA-FL and ChiA domains to immobilized commercially available mucin extracts by ELISA using anti-His antibodies (Fig. 4C). All ChiA samples displayed significant adhesion to mucins isolated from porcine stomachs (type II and III) but this was not observed with a mucin extract from bovine submaxillary glands (type I-S). This confirms that ChiA has additional specificity for non-chitinous ligands and implies that ChiA could mediate the attachment of host glycoproteins to the *Legionella* surface. To assess this, we incubated *L. pneumophila* 130b wild-type and NU318 (*chiA*) mutant strains with type I-S, II and III mucin extracts followed by wheat germ agglutinin and measured their binding to the bacterial surface by flow cytometry. Whether examined at 25°C or 37°C, type II and III extracts showed strong association to both *L. pneumophila* strains (Fig. 4D) but submaxillary gland mucins did not (data not shown). However, there was no significant difference in binding of the type II and III mucins between wild-type and *chiA* mutant strains, which indicates that other factors apart from ChiA are present on the *Legionella* surface that can also recognize mucins and in the case of the mutant, compensate for the loss of ChiA.

### ChiA-CTD is a metal-dependent mucinase

We next examined whether secreted ChiA is able to degrade mucins. Porcine stomach type II mucin extract was incubated with supernatants from *L. pneumophila* 130b wild-type and NU318 (*chiA*) mutant strains or a cocktail of enzymes (pepsin, pronase, *β*-N-acyltylglucosamidase, fucosidase) with known activity against mucins, and then analysed by immunoblotting using wheat germ agglutinin (Fig. 5A**, left panel**). While the majority of the mucin extract ran at >500 kDa, after incubation with the mucinase cocktail there was a reduction in high molecular weight species and the appearance of a new band at ∼200 kDa. When the extract was incubated with *L. pneumophila* 130b wild-type supernatant there was again a reduction of high molecular weight material but also addition of a fragment at ∼100 kDa, while the *chiA* mutant supernatant produced a profile that was more similar to the control. When the experiment was performed using a greater amount of mucin, the difference between the wild type and mutant was even more evident, revealing that the presence of the 100 kDa degradation product was dependent upon ChiA (Fig 5A**, right panel**).

We then tested the ability of recombinant ChiA to degrade type II mucin extracts *in vitro* by immunoblotting and compared its profile to that of recombinant SslE mucinase, an *E. coli* M60 family aminopeptidase (32). Incubation of type II mucin with ChiA-FL and SslE resulted in a significant reduction of detectable carbohydrate, showing that both enzymes process mucins into lower molecular weight fragments and confirmed that *L. pneumophila* degradation of mucins is a direct result of ChiA activity. (Fig. 5B). SslE uses a HExxH motif to coordinates Zn^2+^ in its active site (33) and in the presence of the metal chelating agent ethylenediaminetetraacetic acid (EDTA) we could not detect activity for SslE. Likewise, in the presence of EDTA ChiA-FL could not process mucins which demonstrates that the mucinase activity of ChiA is also metal-dependent.

### ChiA-CTD is a novel peptidase

Mucins contain a large number of serine/threonine rich repeat sequences which are highly O-glycosylated (34). Analysis of the type II mucin extract by 1D ^1^H-NMR revealed sharp peaks that originate from partially degraded low molecular weight mucin fragments and we used this to further probe changes in the mucin structure on addition of ChiA. Incubation with ChiA-FL, ChiA-CTD and SslE but not ChiA-NT resulted in a loss of signal in the type II mucin extract at ∼1.2 ppm and between ∼2.5 to ∼2.8 ppm and ∼3.8 to 4.2 ppm, whilst in the presence EDTA no significant changes were observed (Fig. 5C, S8). The degradation profile of the type II mucin extract incubated with ChiA-CTD and SslE were identical and changes in the ^1^H-NMR spectra are localized to the expected chemical shift ranges for non-anomeric carbohydrate ring protons and the amino acids side chain protons of aspartate, asparagine and threonine. This clearly shows that the C-terminal domain of ChiA has dual enzymatic activity and supports ChiA and SslE having similar peptidase mechanisms for the degradation of mucins.

To evaluate this further we performed Molecular Dynamics (MD) simulations and examined the ability of ChiA-CTD to bind Zn^2+^ *in silico*. The protein was ‘soaked’ in a water solution at high Zn^2+^ concentration and the system was then left to evolve over time to identify the regions on the protein surface where Zn^2+^ ions tend to bind. Multiple short simulations were run starting from different random placements of Zn^2+^ ions, for an aggregated simulation time of 1.7 µs. Analysis of the Zn^2+^ spatial distribution function (sdf) calculated on the concatenated trajectories highlighted multiple high Zn^2+^ density sites in the region around the chitinase active site, providing information on the different ways in which Zn^2+^ could bind to the protein in this region. The highest density was found at the chitinase active site (region 1), where Zn^2+^ is coordinated by Asp541, Glu543 and Gln617, with two other sites in close proximity (regions 2,3) coordinated by Glu543 and Q583 (Fig. 6). Binding of zinc in the active site of *Bacillus cereus* ChiNCTU2 has been shown to inhibit chitinase activity (21) and indicates that metal binding could modulate the different enzyme activities in ChiA.

A unique cluster of Zn^2+^ sites was also located away from the chitinase active site, near the MPD-2 ligand site in the ChiA-CTD crystal structure, involving residues Asp504 (region 4), His544 (region 5), Glu543 and Gln595 (region 6), Asn547 (region 7) and His506 (region 8) (Fig. 6). To assess whether this second cluster could coordinate a mucinase active site metal we created D504A, H506A, H544A, N547A and Q595A mutants in ChiA-CTD (Fig. S2). All mutants retained their ability to bind chitin in pull-down experiments (Fig. S9) and bind immobilized type II and III mucin extracts in ELISA assays (Fig. S10). We then incubated these proteins and ChiA-CTD^E543M^ with type II mucin extract and inspected their degradation profiles by ^1^H NMR. Mucin incubated with ChiA-CTD^D504A^, ChiA-CTD^H544A^, ChiA-CTD^N547A^ and ChiA-CTD^Q595A^ all produced identical NMR spectra that showed no degradation, while mucins incubated with ChiA-CTD^H506A^ or ChiA-CTD^E543M^ showed loss of signals at ∼1.2 ppm and between ∼2.5 to ∼2.8 ppm and ∼3.8 to 4.2 ppm equivalent to that of native ChiA-CTD (Fig. 7A). This confirms that Asp504, His544, Asn547 and Gln595 form a unique peptidase active site in ChiA which is independent from the adjacent glycosyl hydrolase active site (Fig. 7B).

**Figure 7:**
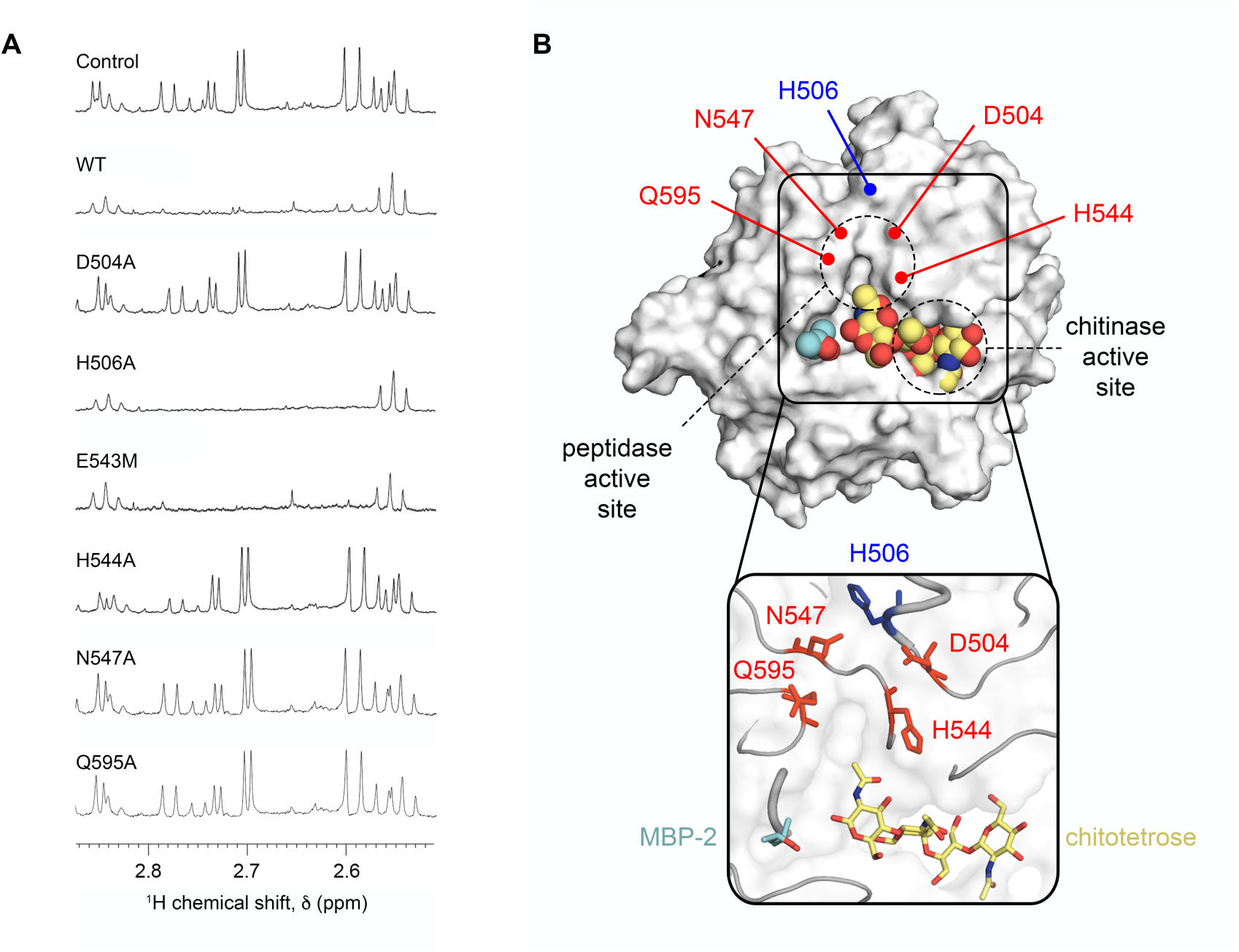
Peptidase active site of ChiA. (A) 1D ^1^H NMR spectra between ∼2.5 to ∼2.9 ppm is shown for type II porcine stomach mucins incubated with ChiA-CTD, ChiA-CTD mutants (D504A, H506A, E543M, H455A, N547A, Q595A) or buffer alone. (B) Surface and cartoon representation of ChiA-CTD bound to MPD-2 (cyan; spheres and sticks) and modelled chitotetrose (yellow; spheres and sticks), highlighting the potential for mucin branched glycan recognition. Residues that form the metal-dependent aminopeptidase active site are highlighted in red.

## Discussion

Chitin is highly abundant in the environment and can function as a source of carbon and nitrogen (35) but several chitinases have been identified as key virulence factors in bacterial disease (31). These include *Enterococcus faecalis* efChiA, *E. coli* ChiA, *Vibrio cholerae* ChiA2, *Francisella tularensis* ChiA, *Listeria monocytogenes* ChiA and ChiB, *Pseudomonas aeruginosa* ChiC, *Salmonela* Typhimurium ChiA and *L. pneumophila* ChiA. Although it is unclear how these enzymes perform these dual functions, there is strong evidence that they interact with host glycoconjugates and through their localization and/or enzymatic activity are able to modulate host defence mechanisms (31). We have determined that *L. pneumophila* ChiA has activity against stomach derived mucins and produces an identical degradation profile to the M60-family *E. coli* zinc-aminopeptidase SslE (36, 37). Recent structural analysis of *Bacteroides thetaiotaomicron* BT4244, *Pseudomonas aeruginosa* IMPa and *Clostridium perfringens* ZmpB M60 proteins has revealed unique structural adaptations that allow them to accommodate different glycan sequences while all cleaving the peptide bond immediately preceding the glycosylated residue (33). Similarly, enterohemorrhagic *E. coli* StcE is an M66-family zinc metalloprotease that recognizes distinct peptide and glycan motifs in mucins and then cleaves the peptide backbone using an extended HExxHxxGxxH motif (38, 39). We have shown that *L. pneumophila* ChiA functions in a similar fashion to SslE but as it does not contain a HEXXH motif, ChiA represents a new class of peptidase that can degrade mammalian mucins via a novel mechanism.

We have identified four residues in ChiA that are essential for peptidase activity (Asp504, His544, Asn547 and Gln595) and likely function by coordinating a Zn^2+^ or other catalytic metal (33). While Asp504 is located in an augmented *α*3 helix, His544, Asn547 and Gln595 are positioned within conserved GH18 secondary structure (Fig. S5). However, sequence alignment of *L. pneumophila* ChiA with other virulent bacterial chitinases, including the mucin degrading *V. cholerae* ChiA2 (40), does not show conservation of these residues and modelling of their tertiary structures using the Phyre2 server (18) also highlights significant differences within their chitin binding surfaces. This implies that other virulent bacterial chitinases either promote pathogenesis using an alternative mechanism or that the specific location of the peptidase active site is unique to each enzyme and shapes their glycan specificity and function.

Mucins are heavily glycosylated cell surface exposed transmembrane proteins or secreted gel-forming proteins of the mucosal barrier that act as a first line of defence against bacterial infection and vary significantly in their structures and glycosylation profiles (41). Mucins derived from the lung are not commercially available but we have shown that ChiA has specificity for and can degrade mucins purified from the porcine stomach. The normal stomach mucosa is characterised by expression of MUC1, MUC5AC, and MUC6 mucins (42), however, MUC1 and MUC5AC are also major mucins expressed in the human airway and lung (43). MUC5AC is composed of T-antigen (Galβ1-3GalNAcαSer/Thr), core 2 (GlcNAcβ1-6(Galβ1-3)GalNAcαSer/Thr) and sialyl T-antigen (NeuAcα2-6(Galβ1-3)GalNAcαSer/Thr) core glycan structures (44) and within the ChiA C-terminal domain the MPD-2 and chitin binding sites have obvious potential to capture these GlcNAc-like linear and branched chains (Fig. 7B). Our data suggests that ChiA can process major mucins expressed in the lung and liberate nutrients and/or facilitate bacterial penetration of the alveolar mucosa.

Secreted ChiA has been detected in bacterial culture supernatants and during *L. pneumophila* infection of cultured amoebae and human macrophages (15, 26). We previously observed that in the early stages of macrophage infection ChiA and another type II substrate ProA are exported to the host cytoplasm and localize to the surface of the LCV (26). The ability of ChiA to bind non-chitin substrates implies that during intracellular infection ChiA interacts with other specific host cytoplasmic glycoproteins and may modulate host pathways either through their requisitioning to the LCV or through their hydrolysis. In this study we have now shown that ChiA can also bind the *L. pneumophila* surface, through its N3 domain, although ProA does not. In other bacteria, type II substrates can associate with their outer membrane via acetylation of their N-terminus, interactions with other outer membrane proteins or through recognition of lipopolysaccharides (27, 28). While the mechanism by which ChiA is targeted to the *Legionella* outer membrane is still unclear, this may be shared during localization of ChiA to the LCV, yet interactions that tether ProA to the LCV are clearly not also present on the *L. pneumophila* surface.

Our examination of recombinant ChiA subdomains has revealed that the N-terminal domains and C-terminal chitinase/peptidase domain can all bind porcine stomach mucins and implies that host mucins can be sequestered and processed on the bacterial surface. However, we did not observe significant differences between mucin binding to *L. pneumophila* 130b wild-type or *chiA* mutant strains. This demonstrates that additional uncharacterized mucin binding proteins are also present on the bacterial surface and indicates that manipulation of the host musosa is an important pathogenic mechanism of *L. pneumophila*. The multifaceted nature of ChiA makes it a highly versatile virulence factor of *L. pneumophila* and likewise a target for controlling *L. pneumophila* infection. As a surface associated protein ChiA is a promising vaccine target and our structural characterization may provide a platform to initiate vaccine development.

## Acknowledgements

KHR, RS and JAG were supported by the MRC (MR/M009920/1). PC was supported by the Academy of Medical Sciences/Wellcome Trust (SBF002/1150). TJP was supported by an EPSRC studentship and BD was supported by a BBSRC studentship. Work in the Cianciotto lab was supported by NIH grant R01AI 043987. LSG and RCW were also partly supported by NIH training grants T32 GM08061 and T32 AI0007476, respectively. We thank the beamline scientists at I02, I04 and B21 of the Diamond Light Source. This work was supported by the Francis Crick Institute through provision of access to the MRC Biomedical NMR Centre. The Francis Crick Institute receives its core funding from Cancer Research UK (FC001029), the UK Medical Research Council (FC001029), and the Wellcome Trust (FC001029). MD simulations were performed using time on ARCHER granted via HECBioSim, supported by EPSRC (EP/L000253/1).

## Author Contributions

Conceived and designed the experiments: KHR, LSG, PC, RW, AF, NPC, JAG. Performed the experiments: KHR, LSG, PC, RCW, RS, TP, BD, AF, JAG. Analyzed the data: KHR, LSG, PC, RCW, AF, NPC, JAG. Contributed reagents/materials/analysis tools: AF, NPC, JAG. Wrote the paper: AF, NPC, JAG.

---

**Figure S1:**
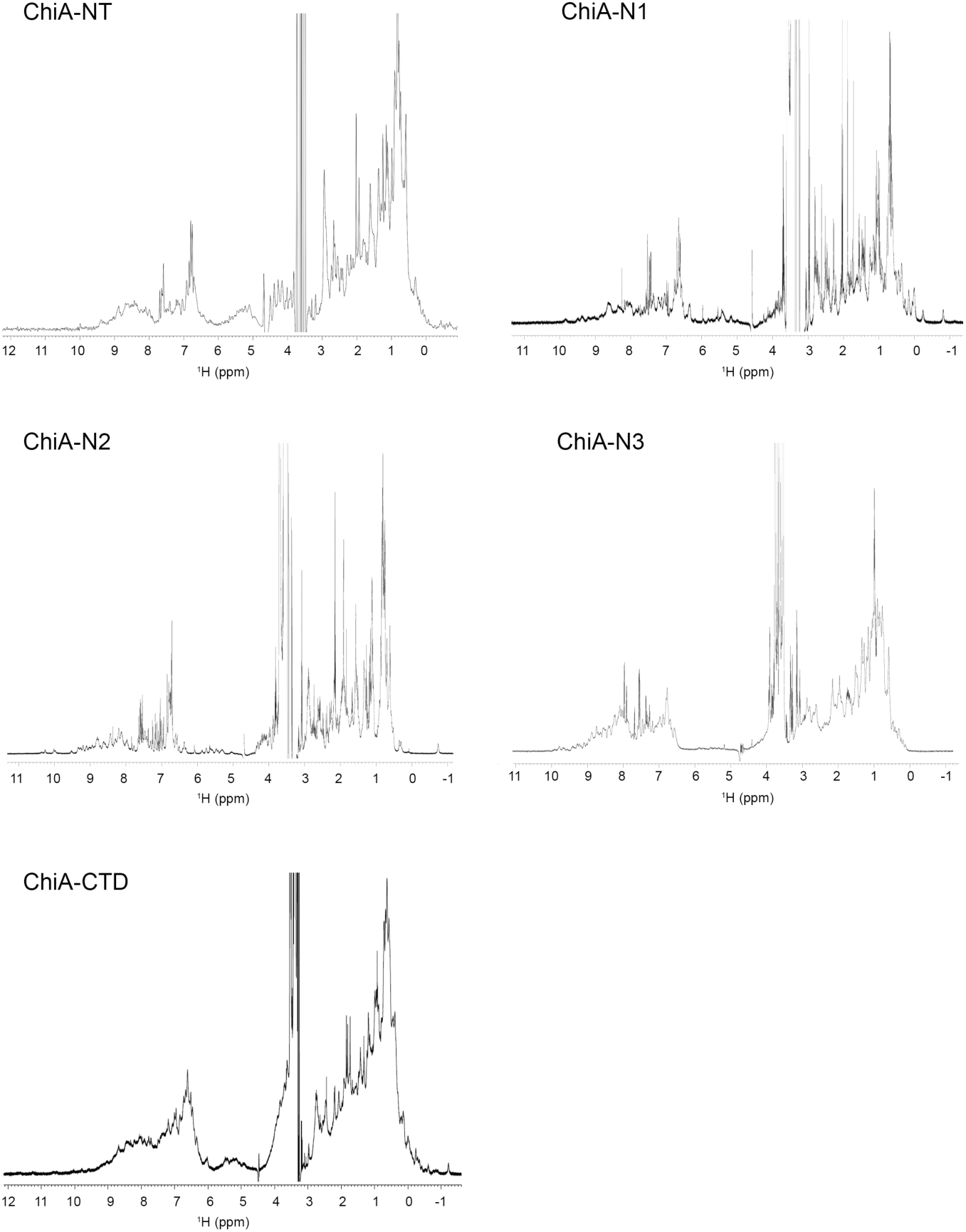
1D ^1^H NMR spectra of ChiA subdomains. The methyl region of the NMR spectra includes high-field proton resonances observed at low chemical shifts (<0.5 ppm), which indicate the presence of characteristic clusters of aromatic and methyl groups in the core of a structured protein. In addition, the envelope of peaks resonating at high chemical shift (>8.5 ppm) correspond to highly ordered backbone amides present in secondary structure elements.

**Figure S2:**
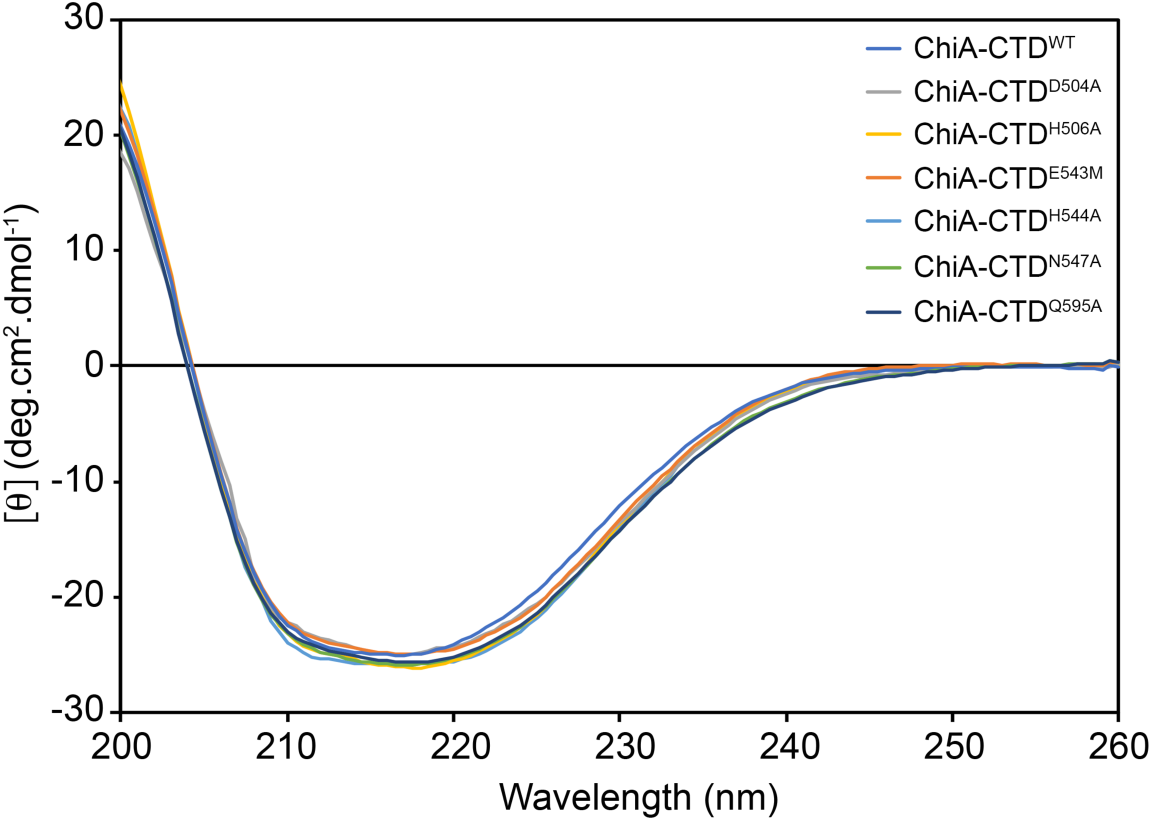
Circular dichroism (CD) spectra of ChiA-CTD constructs. The negative bands between ∼210 to ∼ 220 nm and positive band at 200 nm is indicative of a mixed *α*/*β* protein fold. The spectra for wild type (WT) ChiA-CTD and mutants are in essence identical and demonstrates that these mutations do not perturb the structure of the CTD domain.

**Figure S3:**
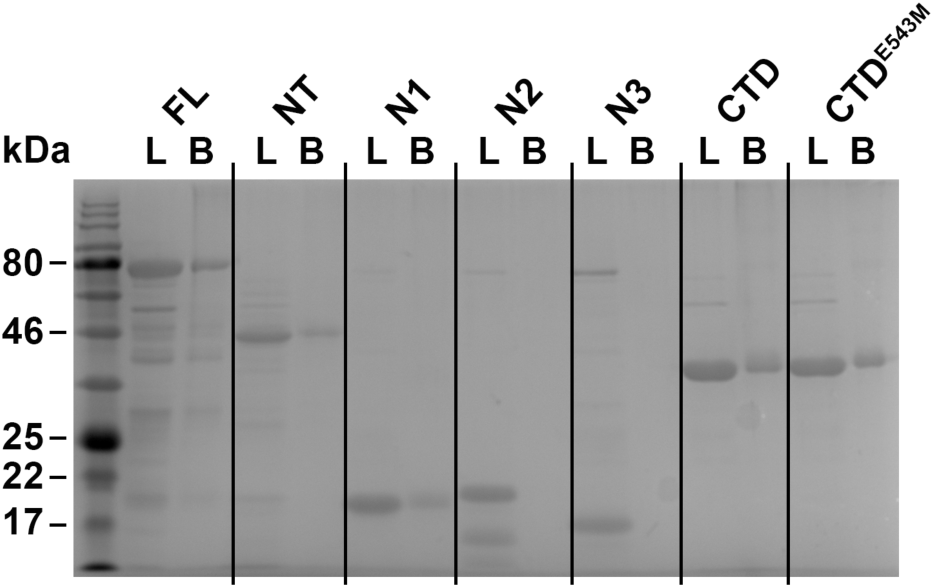
Chitin-resin pull down with ChiA fragments. SDS-PAGE gels loaded with ChiA and subdomains or BSA control either before incubation with chitin beads (L) or after elution from the beads (B).

**Figure S4:**
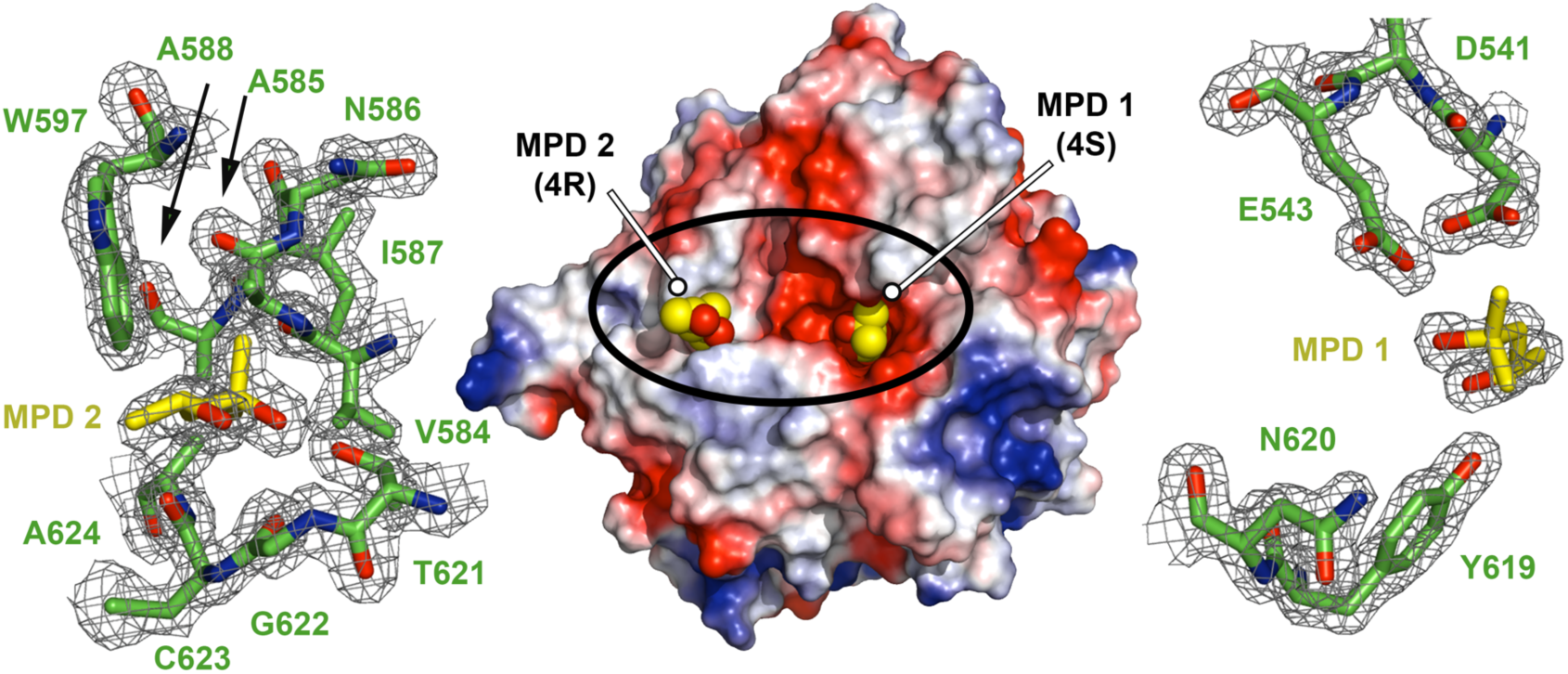
MPD bound to ChiA-CTD. Electrostatic surface potential representation of ChiA-CTD with two molecules of MPD shown as spheres (MPD1: 4S enantiomer; MPD2: 4R enantiomer). Each binding site is expanded and the *σ*A weighted electron density maps contoured at 1.0 r.m.s. are shown.

**Figure S5:**
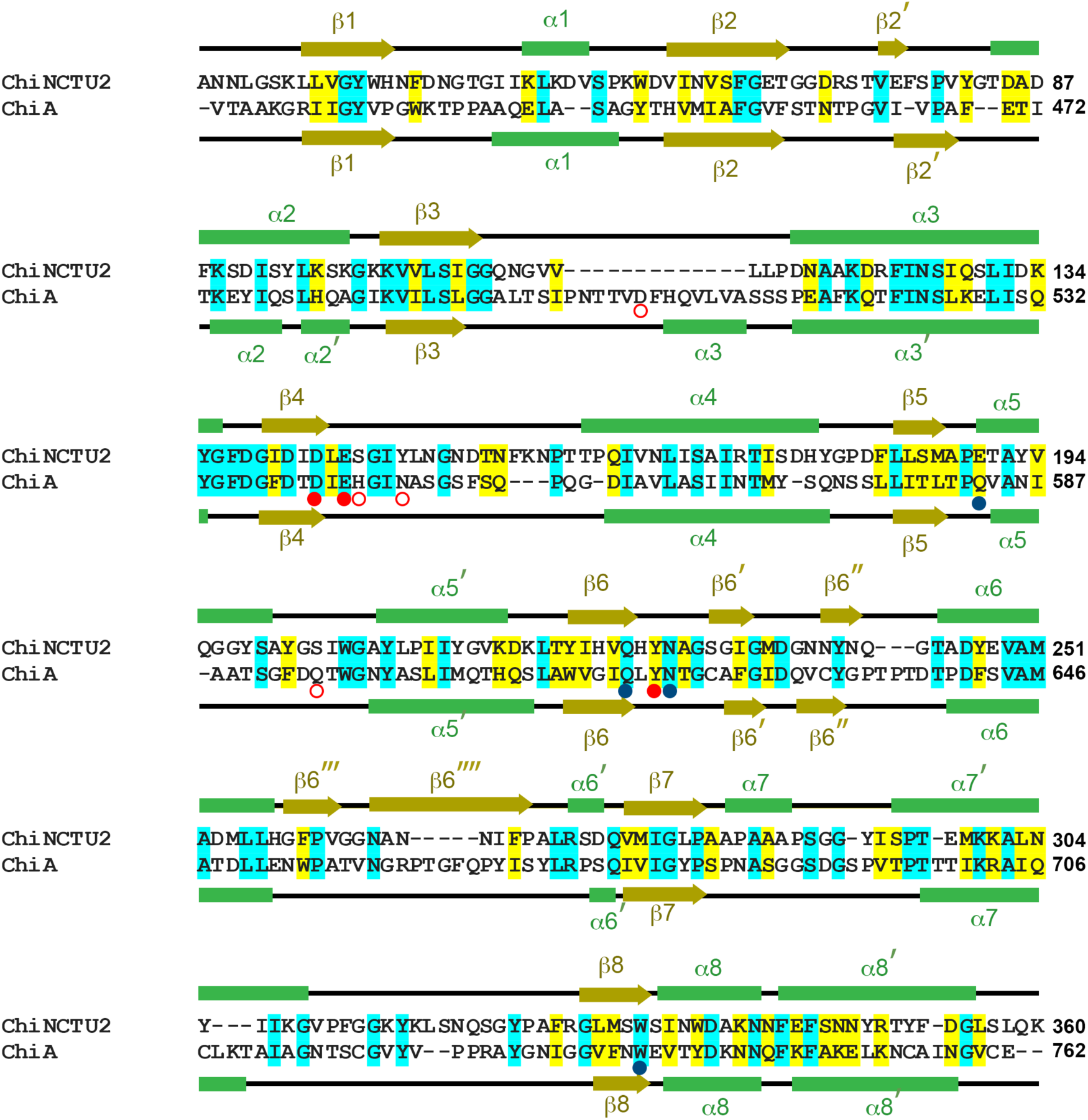
Sequence alignment of *Legionella pneumophila* ChiA-CTD and *Bacillus cereus* ChiNCTU2. Secondary structure elements of ChiNCTU2 and ChiA-CTD are shown above and below, respectively (green rectangle: *α*-helix; gold arrow: *β*-strand). Amino acid identities and similar residues are indicated by background shading in cyan and yellow, respectively. Catalytic chitinase residues and chitin binding residues in ChiNCTU2 are indicated with red and blue filled circles, respectively. Mucinase active site residues in ChiA-CTD are shown as open red circles.

**Figure S6:**
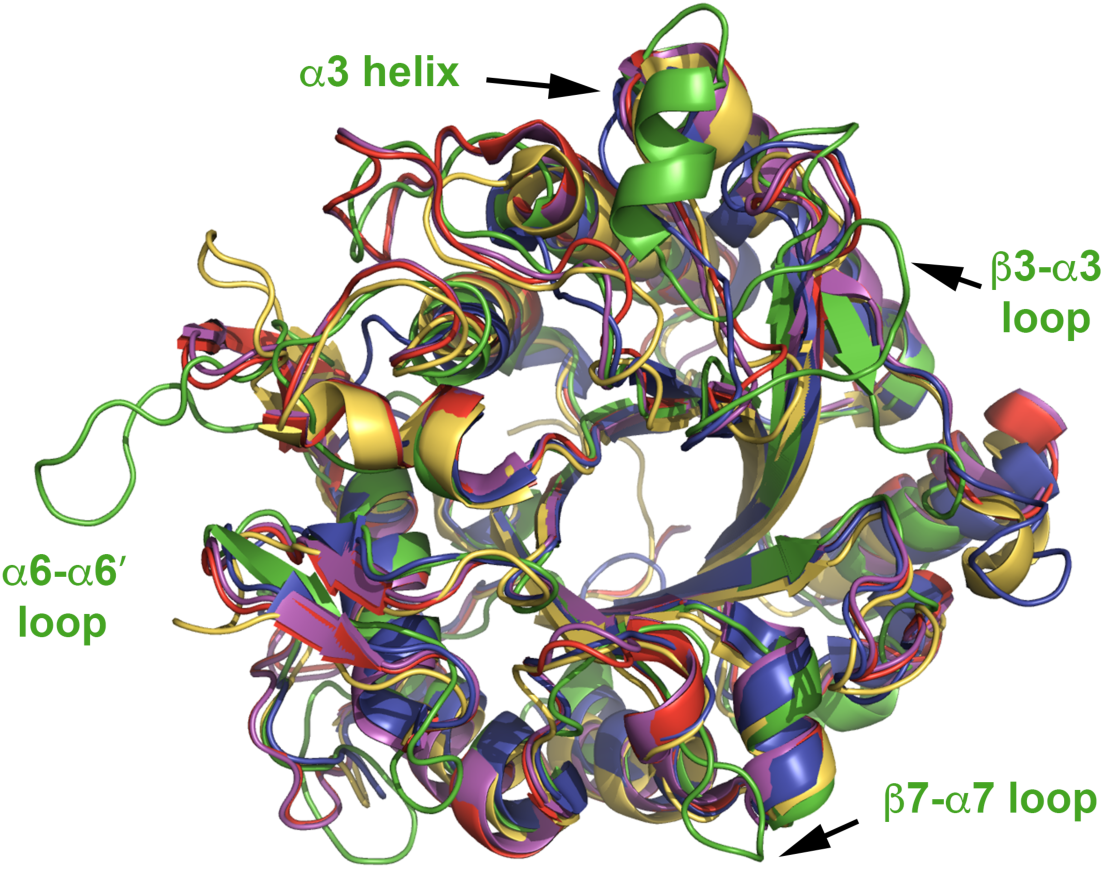
Superposition of ChiA-CTD tertiary homologs. *L. pneumophila* ChiA-CTD is green, *Bacillus cereus* ChiNCTU2 is purple (PDB ID code 3n18) (45), *Bacillus anthracis* Chi36 is red (PDB ID code 5kz6, *Chromobacterium violaceum* ChiA is yellow (PDB ID code 4tx8) and *Streptomyces coelicolor* ChiA is blue (PDB ID code 3ebv). Augmented loop and helical structures in *L. pneumophila* ChiA-CTD are annotated.

**Figure S7:**
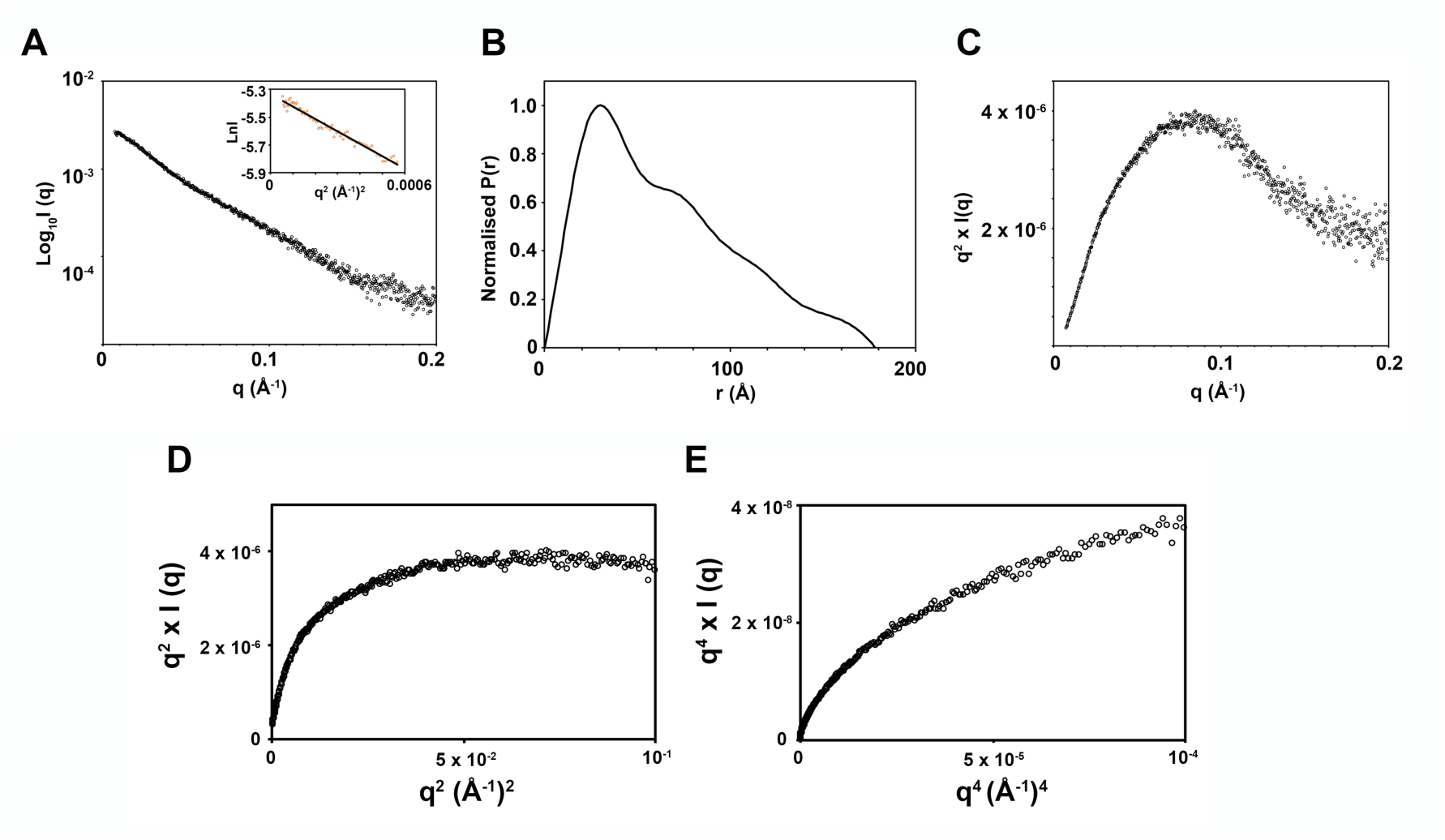
SAXS analysis of ChiA-FL. (A) Experimental scattering curve of ChiA-FL (black open circles). Inset: Guinier Region (orange open circles) and linear regression (black line) for Rg evaluation. (B) Shape distribution [P(r)] function derived from SAXS analysis for ChiA. (C) Kratky, (D) Kratky-Debye and (E) Porod-Debye plots indicate that ChiA is a highly dynamic particle in solution.

**Figure S8.**
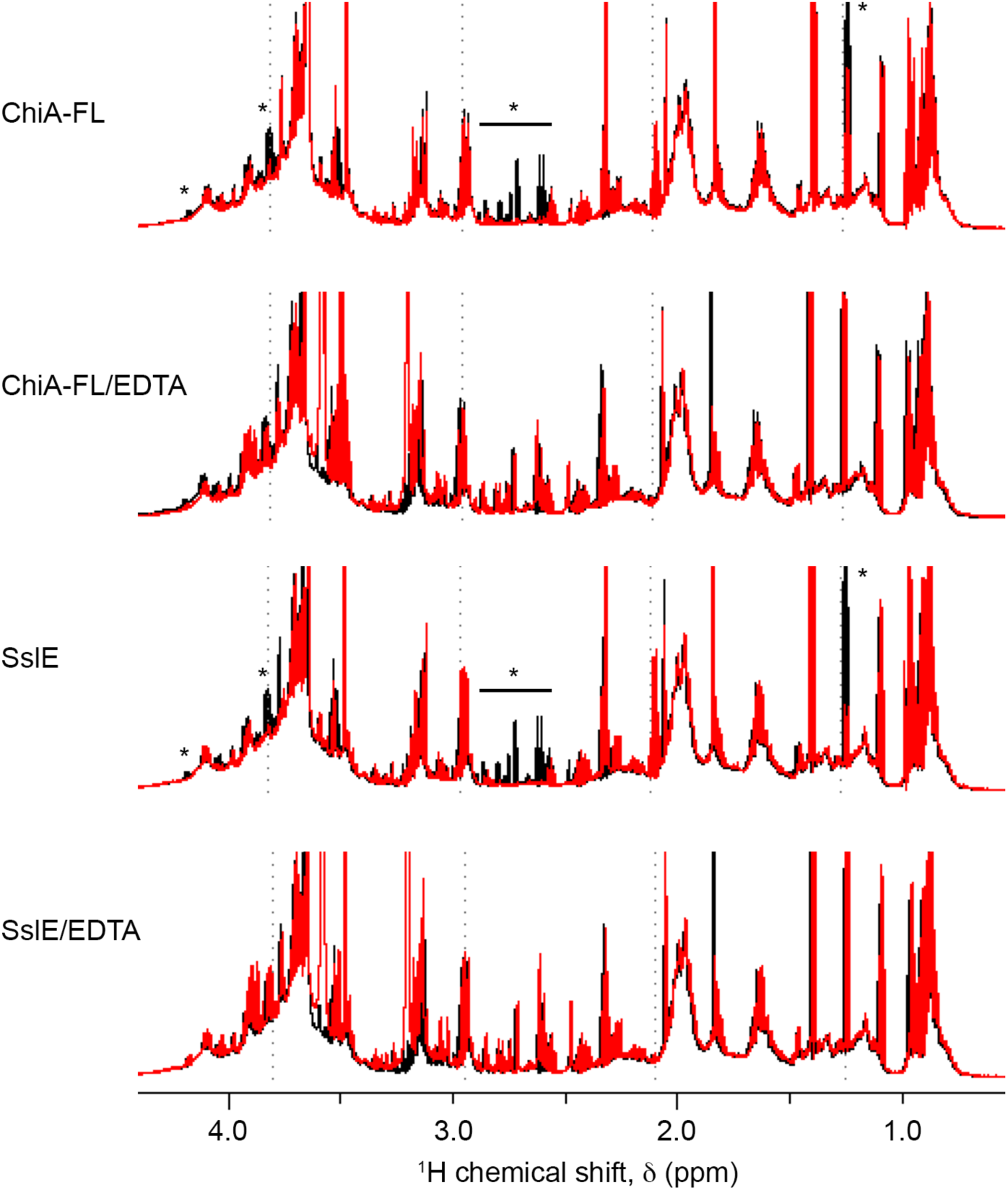
1D ^1^H NMR spectra showing mucinase activity of ChiA. Type II mucin extract incubated at 24°C for 12 hrs (black) is compared to type II mucin extract with either full length ChiA (ChiA-FL), ChiA-FL and EDTA, SslE or SslE and EDTA incubated at 24°C for 12 hrs (red). The region between ∼0.5 and 4.5 ppm is shown and significant differences between the spectra are highlighted with an asterisk.

**Figure S9:**
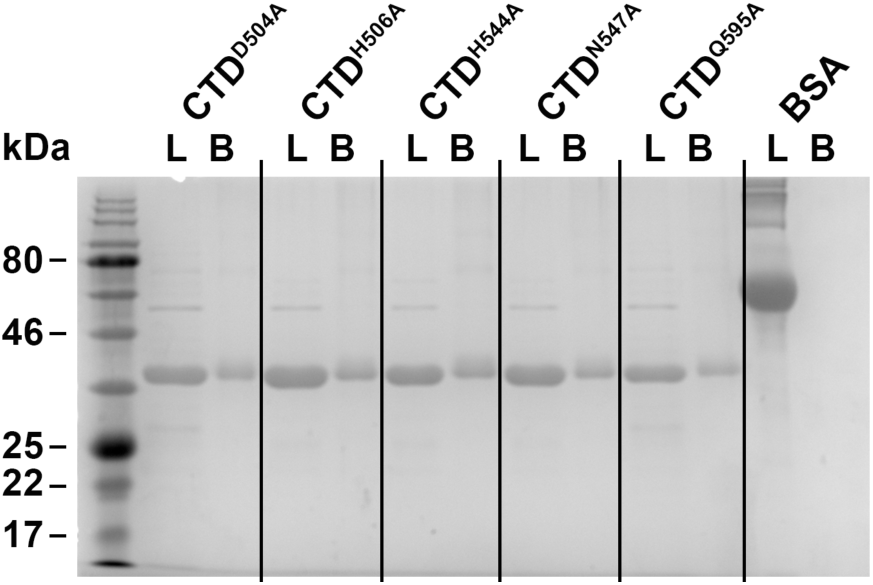
Chitin-resin pull down with ChiA mutants. SDS-PAGE gels loaded with ChiA-CTD mutants or BSA control either before incubation with chitin beads (L) or after elution from the beads (B).

**Figure S10:**
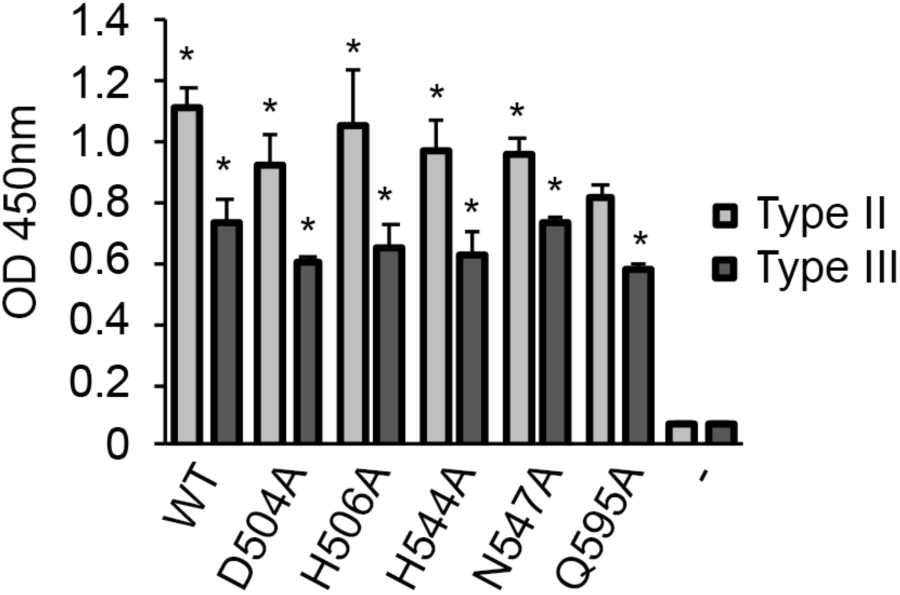
Mucin binding of ChiA-CTD mutants. (A) ELISA analysis of binding between immobilised type II or III mucin extracts and His-tagged wild-type ChiA-CTD (WT) and ChiA-CTD mutants (D504A, H506A, H544A, N547A, Q595A). Anti-His-tag antibody conjugated to HRP was used to measure OD_450 nm_ values. BSA-coated wells were used as controls. Data represent the mean and standard deviation for triplicate experiments. *, *P* < 0.001; verses control empty well by two-tailed Student’s test.

## Supplementary Tables

**Table S1:** Tertiary structure predictions of ChiA N-terminal subdomains.

| ChiA domain | Protein Name | Organism | Function | Sequence identity | PDB code | Ref | Prediction server |
| --- | --- | --- | --- | --- | --- | --- | --- |
| N1 | RsgI2 | *Clostridium thermocellum* | Putative cellulosomal family 3 carbohydrate binding module | 16% | 4B9P | (46) | Phyre2/Robetta |
| N1 | ScaA | *Acetivibrio Cellulolyticus* | Family 3B carbohydrate binding module – binds strongly to cellulose | 21% | 3ZQW | (47) | Phyre2 |
| N1 | CipC | *Clostridium cellulolyticum* | Family 3A carbohydrate binding module – binds strongly to cellulose | 20% | 1G43 | (48) | Phyre2 |
| N1 | Endo-glucanase D | *Clostridium cellulovorans* | C-terminal carbohydrate binding module | 17% | 3NDY | N/A | Phyre2 |
| N1 | Chi18aC | *Streptomyces coelicolor* | Chitinase chitin binding domain | 18% | 2RTT | N/A | Phyre2 |
| N1 | RsgI3 | *Clostridium thermocellum* | Putative cellulosomal family 3 carbohydrate binding module | 14% | 4B97 | (46) | Phyre2 |
| N1 | CBDcex | *Cellulomonas fimi* | Cellulose carbohydrate binding module | 19% | 1EXH | (49) | Phyre2 |
| N2 | RG-lyase | *Aspergillus aculeatus* | Rhamnogalacturonan lyase fibronectin type III domain | 19% | 1NKG | (50) | Phyre2 |
| N2 | CbhA | *Clostridium thermocellum* | Cellobiohydrolase A fibronectin type III domain | 16% | 3PE9/3PDG | (51) | Phyre2 |
| N2 | Tenascin | *Chicken* | Tandem fibronectin type III domains | 12% | 1QR4 | (52) | Robetta |
| N3 | ChiA | *Autographa californica nuclear polyhedrosis virus* | N-terminal chitinase domain – chitin binding | 15% | 5DF0 | N/A | Phyre2 |

**Table S2:** Diffraction data and refinement statistics.

|  | Native | Iodide |
| --- | --- | --- |
| Crystal Parameters |  |  |
| Space group | *C*2 | *C*2 |
| Cell dimensions (Å; °) | *a*=97.37, *b*=56.64, *c*=128.96; β=93.79 | *a*=97.25, *b*=57.35, *c*=129.42; β=93.43 |
| Data collection |  |  |
| Beamline | DLS I04 | DLS I02 |
| Wavelength (Å) | 0.97949 | 1.70000 |
| Resolution (Å) | 64.34-1.71<br>(1.75-1.71) | 64.59-1.89<br>(1.94-1.89) |
| Unique observations | 75905 (5555) | 52043 (2309) |
| *R*<sub>sym</sub> | 0.061 (0.658) | 0.189 (3.697) |
| *<I>/σ* | 19.2 (2.8) | 9.7 (1.4) |
| Completeness (%) | 99.9 (99.9) | 91.2 (55.0) |
| Redundancy | 6.8 (6.9) | 13.1 (11.4) |
| Phasing |  |  |
| Figure of merit (acentric/centric) | 0.159/0.139 | - |
| Phasing power isotropic (acentric/centric) | 1.059/0.919 | - |
| Phasing power anomalous | 0.887 | - |
| Refinement |  |  |
| *R*<sub>work</sub>/*R*<sub>free</sub> (%) | 14.5/16.7 | - |
| Number of protein residues | 678 | - |
| Number of water molecules | 669 | - |
| Number of ligands | 2 MPD, 2 MRD | - |
| rmsd stereochemistry |  |  |
| Bond lengths (Å) | 0.012 | - |
| Bond angles (°) | 1.534 | - |
| Ramachandran analysis |  |  |
| Residues in outlier regions (%) | 0.0 | - |
| Residues in favoured regions (%) | 96.6 | - |
| Residues in allowed regions (%) | 100 | - |

Numbers in parentheses refer to the outermost resolution shell.
$R_{\text{sym}} = \sum |I - \langle I \rangle| / \sum I$ where *I* is the integrated intensity of a given reflection and *<I>* is the mean intensity of multiple corresponding symmetry-related reflections.
$R_{\text{work}} = \sum ||F_o| - |F_c|| / \sum F_o$ where *F<sub>o</sub>* and *F<sub>c</sub>* are the observed and calculated structure factors, respectively.
*R*<sub>free</sub> = *R*<sub>work</sub> calculated using 10% random data excluded from the refinement.
rmsd stereochemistry is the deviation from ideal values.
Ramachandran analysis was carried out using Molprobit (53).

**Table S3:** SAXS structural parameters.

|  |  |
| --- | --- |
| SAXS data collection |  |
| Beamline | DLS B21 |
| Wavelength (Å) | 1.0 |
| q Range (Å<sup>-1</sup>) | 0.004 to 0.4 |
| Concentration range (mg/mL) | 0.5 to 4 |
| Structural parameters |  |
| I(0) | 5.36e-03 ± 2.6e-05 |
| *R<sub>g</sub>* (nm) (from Guinier) | 5.43 ± 0.05 |
| *R<sub>g</sub>* (nm) (from P(*r*)) | 5.45 ± 0.03 |
| *D*<sub>max</sub> (nm) (from P(*r*)) | 17.77 |
| MW (kDa) (from sequence) | 82.6 |
| MW (kDa) (from SAXS) | 89.2 |

**Table S4:** SAXS ensemble optimization parameters.

|  | Run 1 | Run 2 | Run 3 | Average |
| --- | --- | --- | --- | --- |
| $R_{flex}$ (random) | 90.47 (89.14) | 91.60 (89.70) | 92.14 (89.72) | 91.4 (89.5) |
| $R_{sig}$ | 1.22 | 1.22 | 1.25 | 1.23 |
| $\chi^2$ | 1.07 | 1.05 | 1.14 | 1.09 |
| Cluster 1 (%) | 17.2 | 19.3 | 18.2 | 18.2 |
| Cluster 2 (%) | 49.4 | 40.6 | 46.4 | 45.5 |
| Cluster 3 (%) | 33.4 | 40.1 | 35.4 | 36.3 |
| Final ensemble $R_g$ (nm) | 5.40 | 5.43 | 5.39 | 5.41 |
| Final ensemble $D_{max}$ (nm) | 17.11 | 17.28 | 16.73 | 17.04 |

**Table S5:**
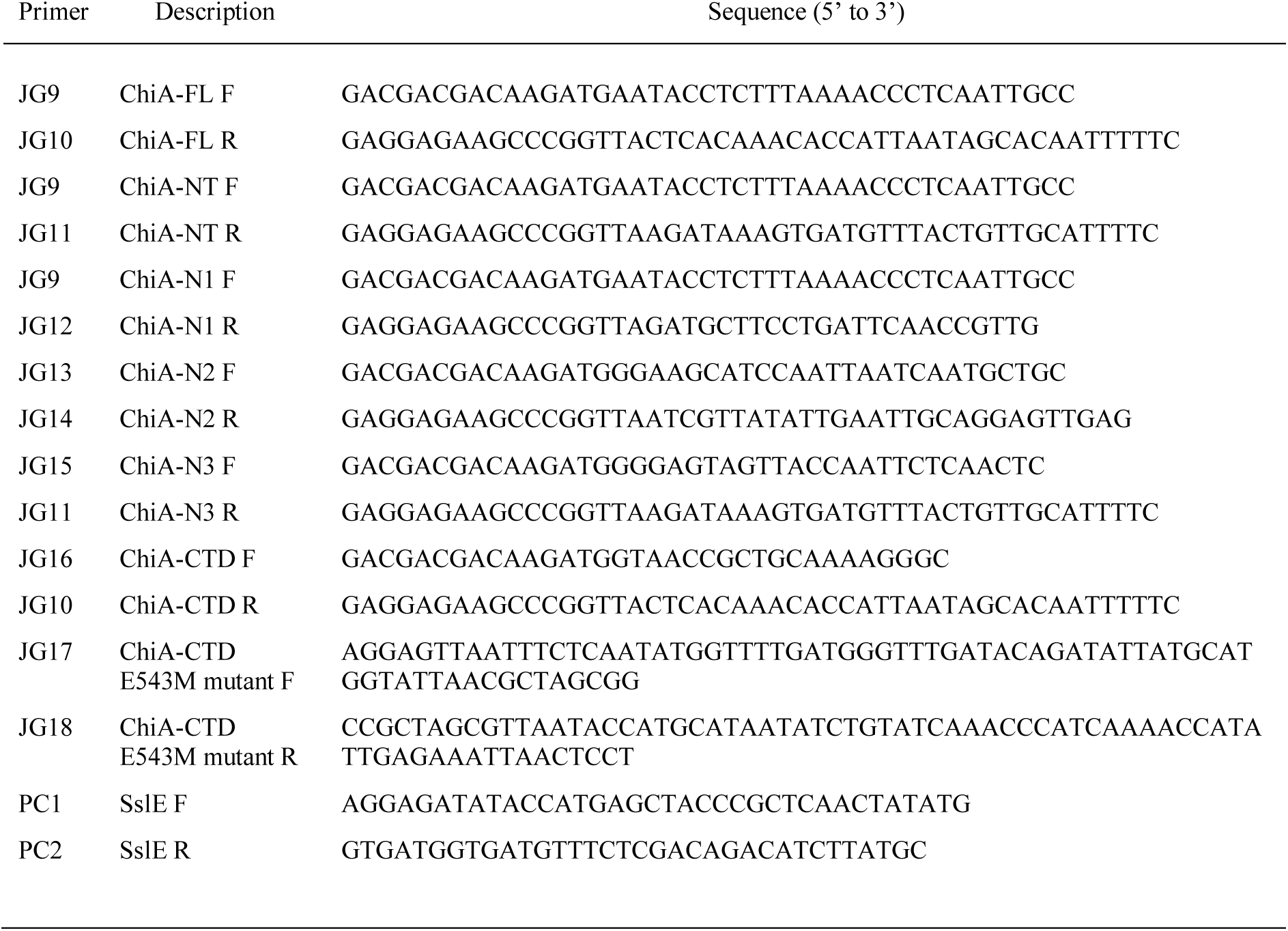
Primers used in this study.

**Table S6:**
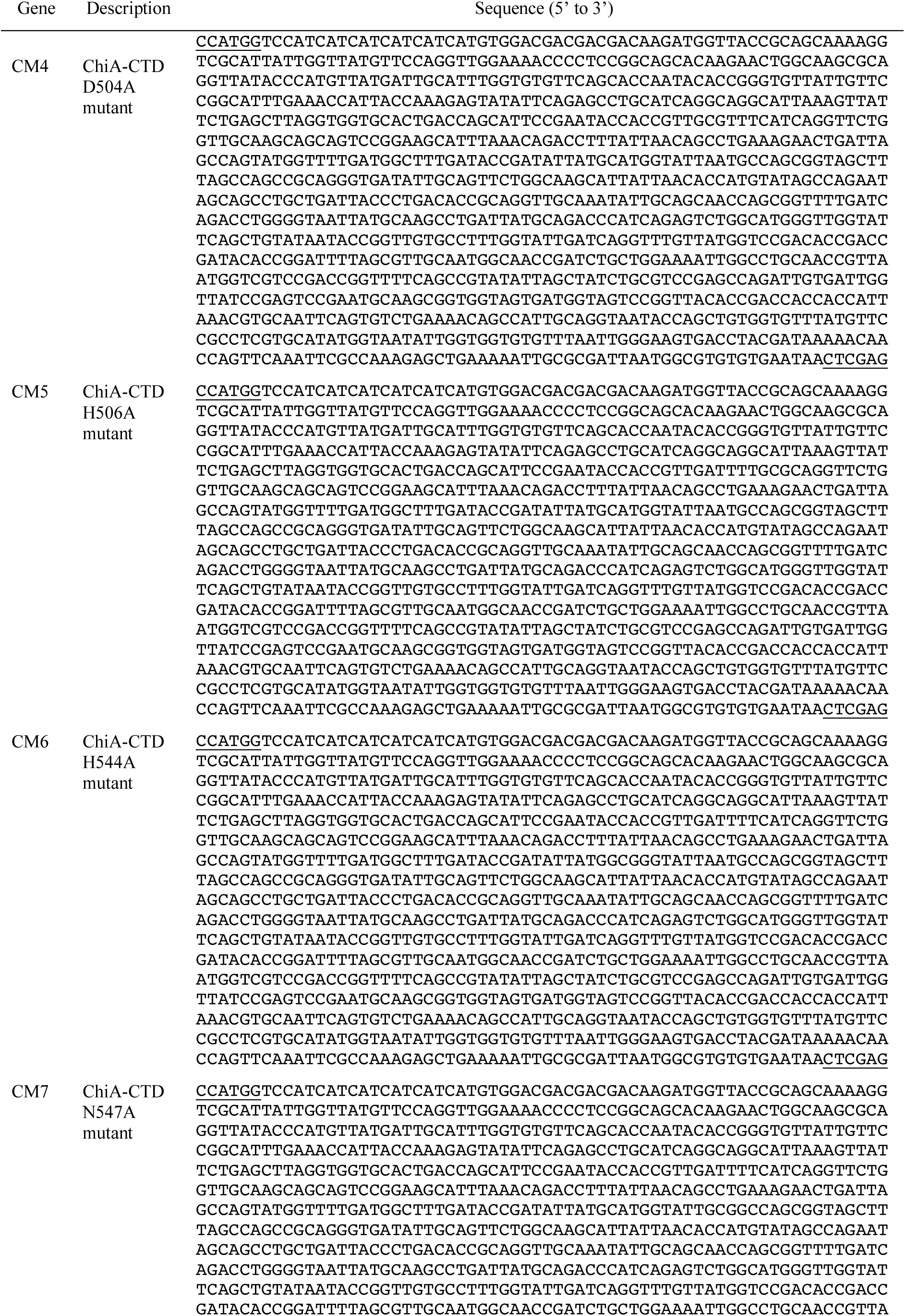
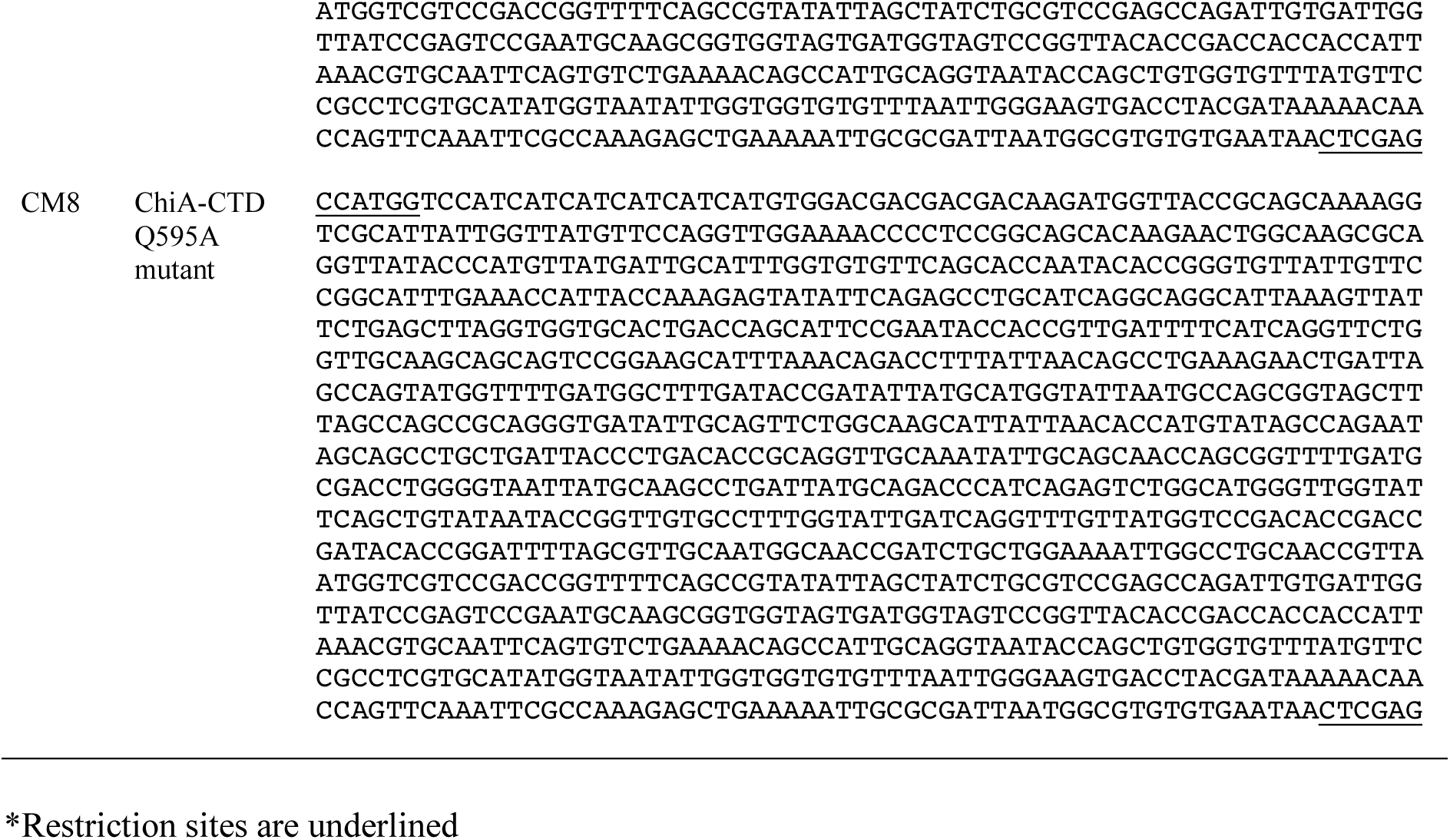
Synthetic genes.

## Methods

### Cloning, expression and purification

Full-length ChiA (ChiA-FL; residues 1–762), minus the N-terminal periplasmic signal sequence, the ChiA N-terminal region (ChiA-NT; residues 1-417), the ChiA N1-domain (ChiA-N1; residues 1-140), the ChiA N2-domain (ChiA-N2; residues 138-299), the ChiA N3-domain (ChiA-N3; residues 285-417) and the ChiA C-domain (ChiA-CTD; residues 419-762) were amplified from the genomic DNA of *L. pneumophila* strain 130b and cloned into the N-terminal His_6_-tagged vector pET-46 Ek/LIC (Table S5). SslE (residues 90-1520), minus the N-terminal periplasmic signal sequence and mature SslE N-terminal proline-rich region, were amplified from the genomic DNA of *E. coli* strain W and cloned into the C-terminal His_6_-tagged vector pOPINE (Table S5). These were transformed into *E. coli* SHuffle cells (New England Biolabs) and grown at 37°C in LB media with 100 µg/ml ampicillin. Expression was induced with 1 mM isopropyl-d-1-thiogalactopyranoside (IPTG) at an OD_600nm_ of 0.6 and cells were harvested after growth overnight at 18°C. Cells were resuspended in 20 mM Tris–HCl pH 8, 200 mM NaCl, 5 mM MgCl_2_, 1 mg/ml DNase I, 5 mg/ml lysozyme, lysed by sonication and purified using nickel affinity chromatography. All samples were then gel filtered using a Superdex 200 column (GE Healthcare) equilibrated in 20 mM Tris–HCl pH 8, 200 mM NaCl.

### Nuclear magnetic resonance spectroscopy

One-dimensional proton NMR experiments were performed at 25°C on 300 µM ChiA-NT, ChiA-N1, ChiA-N2, ChiA-N3 and ChiA-CTD samples in a buffer containing 20 mM Tris-HCl pH 8.0, 100 mM NaCl, 10% D_2_O. Spectra were recorded on a Bruker Avance III 700 MHz (ChiA-N1, ChiA-N2, ChiA-N3, ChiA-CTD) or 800 MHz (ChiA-NT) spectrometer equipped with cryoprobes and processed within TopSpin (Bruker).

### Site-directed mutagenesis

E543M *chiA-CTD* mutant was created using pET46*chiA-CTD* template DNA with a QuikChange II Site-Directed Mutagenesis Kit (Stratagene) (Table S5). D504A, H506A, H544A, N547A and Q595A *chiA-CTD* mutants were synthesized by Synbio Technologies and cloned into pET28b vector using NcoI and XhoI restriction sites (Table S6). All resulting clones were verified by DNA sequencing and then expressed and purified as described for wild-type ChiA-CTD.

### Circular dichroism

Far-UV CD spectra were measured in a Chirascan (Applied Photophysics) spectropolarimeter thermostated at 20°C. Spectra for wild-type ChiA-CTD and ChiA-CTD^E543M^, ChiA-CTD^D504A^, ChiA-CTD^H506A^, ChiA-CTD^H544A^, ChiA-CTD^N547A^ and ChiA-CTD^Q595A^ mutants (0.05 mg/ml) in 10 mM Tris-HCl pH 8.0, 100 mM NaCl were recorded from 260 to 200 nm, at 0.5 nm intervals, 1 nm bandwidth, and a scan speed of 10 nm/min. Three accumulations were averaged for each spectrum.

### Chitinase activity assay

Enzyme activity was determined using 4-Nitrophenol *b*-D-N,N’,N’’-triacetylchitotriose (Sigma) as a substrate. 10 μl of ChiA-FL, ChiA-NT, ChiA-CTD, ChiA-CTD^E543M^ at 10 μg/ml in PBS were incubated with 90 μl of substrate at 0.4 mg/ml dissolved in Sigma buffer (A4855, pH 5.0) (Sigma) at 37°C for 30 min. The reaction was quenched by adding 200 μl of 100 mM sodium carbonate. Absorbance at 405 nm was measured and corrected for absorption in a control sample with added PBS instead of protein.

### Chitin binding assay

250 ml ChiA-FL, ChiA-NT, ChiA-N1, ChiA-N2, ChiA-N3, ChiA-CTD, ChiA-CTD^D504A^, ChiA-CTD^H506A^, ChiA-CTD^E543M^, ChiA-CTD^H544A^, ChiA-CTD^N547A^, ChiA-CTD^Q595A^ and BSA (Sigma) at 10 μM in 20 mM Tris-HCl pH 8.0, 200 mM NaCl were incubated with 50 ml chitin-resin (Sigma) and incubated whilst shaking for 30 min. The resin was washed three times with 500 μl of the same buffer and then proteins were eluted by incubating the resin in 250 μl of 8 M urea, 1% (w/v) SDS for 30 min whilst shaking. Protein samples prior to incubation with chitin-resin and the eluted protein/chitin-resin slurry were then analysed with SDS-PAGE.

### Crystal structure determination

Crystallization of ChiA CTD-domain (30 mg/ml) was performed using the sitting-drop vapour-diffusion method grown in 0.2 M ammonium acetate, 0.1 M Bis-Tris pH 5.5, 45% (v/v) 2-Methyl-2,4-pentanediol at 293K. Native crystals were flash cooled in liquid nitrogen and diffraction data were collected at 100K on beamline I04 at the Diamond Light Source (DLS), UK. Crystals were also soaked for 1 min in well solution containing 1.0 M NaI, flash cooled in liquid nitrogen and data were collected at 100K on beamline I02 at the Diamond Light Source (DLS), UK. Data were processed with XDS (54) and scaled with AIMLESS (55) using the XIA2 pipeline (56). The structure of ChiA CTD-domain was determined with I-SIRAS. Twenty-one iodide sites were located in ChiA C-domain using SHELXD (57), and then phases were calculated using autoSHARP (58). After automated model building with ARP/wARP (59), the remaining structure was manually built within Coot (60). Refinement was carried out with REFMAC (61) using non-crystallographic symmetry (NCS) and translation-libration-screw (TLS) groups (62, 63), and 5% of the reflections were omitted for cross-validation. Processing and refinement statistics of the final model can be found in Table S2. The coordinates and structure factors have been deposited in the PDB under ID code 6s2x.

### SAXS data collection and analysis

SAXS data were collected on beamline B21 at the Diamond Light Source (DLS), UK at 20 °C. Full-length ChiA in 20 mM Tris–HCl pH 8, 200 mM NaCl were measured at 4, 2, 1 and 0.5 mg/ml concentrations, after gel filtration using a Superdex 200 column (GE Healthcare), over a momentum transfer range of 0.004<*q*<0.4 Å^−1^. A fresh sample of BSA was measured as a standard. Buffer subtraction, intensity normalization, and data merging for the different sample concentrations were performed in SCATTER (DLS, UK). ChiA data collected above 1 mg/ml showed signs of aggregation and were discarded. Further analysis was carried out with the 1 mg/ml data using a q range 0.008<*q*<0.2 Å^−1^. The radius of gyration (Rg) and scattering at zero angle (I(0)) were calculated from the analysis of the Guinier region by AUTORG (64, 65). The distance distribution function (P(r)) was subsequently obtained using GNOM (64, 65), yielding the maximum particle dimension (D_max_). Determination of molecular model ensembles that best fit the SAXS data was performed using EOM2.0 (66, 67). An initial model of ChiA was created from PHYRE2 models of the N1- and N3-domains (residues 7-132; 300-399), a ROBETTA model of the N2-domain (residues 138-290) and our crystal structure of the C-domain (residues 424-762), with domain linker sequences kept unstructured. SAXS structural and EOM parameters can be found in Table S3 and Table S4, respectively.

### ELISA for detection of ChiA on bacterial surface

Bacterial whole-cell ELISA was done as previously described (68), with slight modification. Wild-type *L. pneumophila* strain 130b and isogenic mutants lacking either *chiA* (strain NU318) (69), *mip* (strain NU203) (70), or *proA* (strain AA200) (71) were grown on BCYE agar for 3 days at 37°C. Using a sterile cotton swab, bacteria were resuspended in 1 ml sterile PBS to an OD_660_ 0.3, centrifuged at 10,000 x *g* for 3 min, and then washed once with PBS to remove debris and unbound proteins. Bacteria were fixed in 4% (w/v) paraformaldehyde for 10 min, followed by two 1-ml washes in PBS. Bacteria were resuspended in coating buffer (100 mM bicarbonate/carbonate buffer, pH 9.6) to a final OD_660_ 0.03, and 100 µl of this suspension were added into the wells of Nunc MaxiSorp immunoassay plates (Thermo Fisher Scientific). Following overnight incubation at 4°C, the wells were washed three times with 200 µl of wash buffer (PBS + 0.05% Tween-20), and then 200 µl of blocking buffer (PBS + 0.05% Tween-20 + 5% dried milk) were added for 1 h at 25°C. After removal of the blocking buffer, samples were incubated with 100 µl of primary antibody (i.e., rabbit anti-Mip (72), rabbit anti-ChiA (73), or rabbit anti-ProA (73) diluted 1:10,000 in blocking buffer for 1 h at 25°C. Following three, 200-µl washes with wash buffer, samples were incubated with 100 µl of secondary antibody (anti-rabbit conjugated HRP) diluted 1:1,000 in blocking buffer for 1 h at 25°C. Following five washes with 200 µl wash buffer, samples were incubated with 100 µl 3,3’,5,5’-Tetramethylbenzidine (TMB) substrate for 15 min at 25°C, and then, the reaction was stopped by addition of 50 µl of 2 N sulfuric acid. Absorbance values were measured at 450 nm with wavelength correction of 570 nm using a microplate reader (Synergy H1, BioTek). To confirm that bacterial lysis had not occurred during sample processing and plate coating, *L. pneumophila*-coated wells were probed with an ICDH-specific antiserum that recognizes a cytosolic *L. pneumophila* protein (74), and no signal was detected (data not shown). In order to assess the binding of recombinant ChiA and ChiA subdomains to the *L. pneumophila* surface, *chiA* mutant NU318 was grown on BCYE agar, washed, and resuspended in PBS, as indicated above. Prior to fixation, bacteria resuspended to OD_660_ 0.3 were incubated with 1 µg of recombinant protein in the presence of 1x protease inhibitors (Pierce, Thermo Scientific) in sterile 1.5 ml microcentrifuge tubes with gentle end-over-end mixing for 30 min at 25°C.

Following two 1-ml washes in PBS to remove unbound protein, bacteria were fixed with 4% (w/v) paraformaldehyde and processed for ELISA as described above. Preliminary assays determined that the polyclonal anti-ChiA (73) antiserum was capable of recognizing each of the recombinant ChiA fragments when they were added to wells in the absence of bacteria (data not shown).

### Mucin binding ELISA

Immulon 2-HB 96-well plates (VWR) were coated overnight at 4°C with 50 µl of partially purified mucins from bovine submaxillary glands (type I-S; Sigma) and porcine stomachs (type II and III; Sigma) at 100 µg/ml in 50 mM Carbonate/Bicarbonate pH 9.6. Wells were blocked for 1 hr at 25°C with 200 µl of 0.1% (w/v) bovine serum albumin (BSA) in PBS–0.05% Tween 20 and then washed once with 200 µl of incubation buffer (0.05% (w/v) BSA in PBS–0.05% Tween 20). Wells were then incubated for 3 hrs at 25°C with 50 µl of ChiA-FL, ChiA-NT, ChiA-N1, ChiA-N2, ChiA-N3, ChiA-CTD, ChiA-CTD^D504A^, ChiA-CTD^H506A^, ChiA-CTD^E543M^, ChiA-CTD^H544A^, ChiA-CTD^N547A^ and ChiA-CTD^Q595A^ at 10 µM in incubation buffer. This was followed by four washes with 200 µl of incubation buffer and incubation with 50 µl of anti-His-HRP antibody (Sigma), diluted 1:2000 in incubation buffer for 1 hr at 24°C. After four washes with 200 µl of incubation buffer, 150 µl of *o*-Phenylenediamine dihydrochloride (Sigma) was added for 30 min and then data was recorded at 450 nm.

### Assay for mucin binding to bacteria

*L. pneumophila* wild-type strain 130b and *chiA* mutant NU319, both harbouring a GFP-expressing plasmid (69, 73), were incubated for 3 days on BCYE agar containing IPTG at 1mM. Bacteria were suspended in PBS to an OD of 0.3 (i.e., 1 x 10^9^ CFU per ml), and then 1 ml of the suspension was statically incubated with either 100 µg of type II porcine stomach mucins or type III porcine stomach mucins or PBS alone for 1 h at 25°C or 37°C. Following three washes each consisting of a 1-ml PBS wash solution and a 5-min centrifugation step at 4000 x *g* (for the 37°C samples) or 8000 x *g* (for the 25°C samples), bacteria were incubated with 7.5 µg of Texas Red-tagged wheat germ agglutinin (WGA) for 15 min at 25°C in order to detect mucins bound to the bacteria. After three further washes, as indicated above, the bacteria were resuspended in 1 ml PBS and finally analysed on a BD LSRII flow cytometer using a Texas Red filter and GFP Filter (75).

### Immunoblot for detecting secreted mucinase activity

*L. pneumophila* strains that had been grown for three days on BCYE agar were suspended into 20 ml of BYE broth to an OD_660_ of 0.3 and grown overnight at 37°C to an OD_660_ of 3.0 – 3.3. Bacteria were sub-cultured into fresh BYE medium to an OD_660_ = 0.3 and grown, with shaking, to an OD_660_ of 1.0, which corresponded to the mid-log phase. Supernatants were collected, filtered through a 0.22-µm filter, and concentrated using 10-kDa Amicon concentrators (EMD Millipore UFC901024). 200 µl of concentrated supernatants were incubated with 200 or 400 µg of type II porcine stomach mucins. As controls, the mucins were either incubated in uninoculated BYE broth or in BYE broth containing 50 µl of a known mucinase cocktail, which consisted of 10 µl each of pepsin (0.5 mg/ml), pronase (10 mg/ml), B-N-acyltylglucosamidase (2.5 M), fucosidase (5 U/ml), and DTT (1M) dissolved in 940 µl of ddH_2_0. The various samples were incubated statically for 3 h at 25°C and then subjected to electrophoresis prior to immunoblotting (76). Specifically, reactions were stopped by adding 200 µl of 2x Laemmli buffer and incubating for 5 min at 100°C, and 35 µl of each sample was electrophoresed through a Criterion 4-20% SDS-PAGE gel (Biorad 5671094) for 1.5 h at 250 volts, i.e., until the 55-kDa MW marker migrated to the bottom of the gel. The separated reaction products were transferred onto PVDF membrane over the course of 13 min using the semi-dry Invitrogen Power-Blotter and Power Blotter transfer blotting solution. Following incubation in 1% BSA in TBST for 1 h at room temperature, the membranes were incubated overnight at 4°C with biotinylated wheat germ agglutinin that had been diluted 1:2000 (from a 1 mg/ml stock) in TBST with BSA. After three, 5-min washes with TBST buffer, the membranes were further incubated for 1 h at 37°C with Aviden-HRP that had been diluted 1:2000 in BSA-containing TBST. Finally, subsequent to a series of washes, the blot was incubated for 1 min in 2 ml Amersham ECL reagent and then exposed to X-ray film.

### Immunoblot for detecting recombinant ChiA mucinase activity

Partially purified type II porcine stomach mucins (Sigma) at 0.04% (w/v) in PBS were incubated for 24 hrs at 25°C with 10 μM ChiA-FL (with and without 5 mM EDTA), *E. coli* Ssle (with and without 5 mM EDTA) or PBS as a negative control. These were then run on a 1% (w/v) agarose, 0.1% (w/v) SDS gel at 23V in TAE buffer containing 0.1% (w/v) SDS for 14 hrs and then transferred onto a PVDF membrane by blotting. The membrane was blocked for 1hr with 0.1% (w/v) bovine serum albumin (BSA) in PBS–0.05% Tween 20 at 25°C, followed by the addition of 1 μg/ml biotinylated wheatgerm lectin agglutinin for 2 hrs at 25°C. This was then incubated with avidin-HRP (1:2000 dilution) for 1 hr at 25°C and then treated with enhanced chemiluminescence substrate (ECL; Pierce) before detection by enhanced chemiluminescence.

### Detection of recombinant ChiA mucinase activity by NMR

Type II porcine stomach mucins (Sigma) at 0.5% (w/v) in PBS-10% D_2_O were incubated for 12 hrs at 25°C with 10 μM ChiA-FL (with and without 5 mM EDTA), ChiA-NT, ChiA-CTD (with and without 5 mM EDTA), ChiA-CTD^E543M^, ChiA-CTD^D504A^, ChiA-CTD^H506A^, ChiA-CTD^H544A^, ChiA-CTD^N547A^, ChiA-CTD^Q595A^, *E. coli* SslE (with and without 5 mM EDTA) or PBS a negative control. Samples were then analysed by NMR at 25°C using a Bruker Avance III 700 MHz spectrometer equipped with cryoprobe.

### Molecular Dynamics

MD simulations and analyses were performed using GROMACS 2016 v3 (77) using a protocol similar to Ref (78). The protein was described using the Amber99SB*-ILDN force field (79) and solvated using a truncated octahedral box of TIP3P water molecules. A minimal distance of 12 Å was set between the protein and the walls of the box. The charge of the ionisable residues was set to that of their standard protonation state at pH 7. Zn^2+^ions were added by randomly replacing water molecules. A high Zn^2+^concentration (0.75 M) was used to have a faster sampling of possible Zn^2+^ sites around the protein surface. Cl^-^ counterions were added to neutralise the system.

Periodic boundary conditions were applied. The equations of motion were integrated using the leap-frog method with a 2-fs time step. The LINCS (80) algorithm was chosen to constrain all covalent bonds in the protein, while SETTLE (81) was used for water molecules. The Particle Mesh Ewald (PME) (82) method was used for electrostatic interactions, with a 9-Å cutoff for the direct space sums, a 1.2-Å FFT grid spacing, and a 4-order interpolation polynomial for the reciprocal space sums. A 9-Å cutoff was used for van der Waals interactions. Long-range corrections to the dispersion energy were included.

Each system was minimised through 3 stages with 2000 (positional restraints on heavy atoms) + 3000 steps of steepest descent, followed by 2000 steps of conjugate gradient. Positional restraints on heavy atoms were initially set to 4.8 kcal/mol/Å^2^ and they were gradually decreased to 0 in 1.5 ns, while the temperature was increased from 200 to 300 K at constant volume. The system was then allowed to move freely and was subjected to 1-ns equilibration in NVT conditions at T = 300 K. This was followed by a 2-ns equilibration in NPT conditions with T = 300 K and p = 1 bar. For these equilibration steps, the Berendsen (83) algorithm was used for both temperature and pressure regulation with coupling constants of 0.2 and 1 ps, respectively. At last, a 2-ns NPT equilibration was run after switching to the v-rescale thermostat (84) with a coupling constant of 0.1 ps and the Parrinello-Rahman barostat (85) with a coupling constant of 2 ps. Production NPT runs were then performed for 50 ns, saving the coordinates every 1 ps. Multiple replicas (34) were run, with each replica starting from a different configuration of the ions around the protein, for an aggregated simulation time of 1.7 µs (34 × 50 ns).

The spatial distribution function (86) (sdf) of Zn^2+^ around the protein was calculated with the *gmx spatial* tool from GROMACS. Trajectories from the different replicas were first concatenated together and each frame was aligned through a best-fit superposition to the starting frame using the protein coordinates. A 0.5-Å grid spacing was used for the sdf calculation. The average of non-null sdf values was calculated and isosurfaces connecting points with sdf = 20, 25 and 30 x average sdf were considered.

